# Evolutionary landscape of oral microbiome over 100,000 years

**DOI:** 10.1101/2025.11.01.685986

**Authors:** Ling Zhong, Heng Li, Shuting Xia, Shichang Xie, Raffaella Bianucci, Shengkai Li, Sirirak Supa-Amornkul, Camilla Speller, Qing Shi, Yinqiu Cui, Huilin Yang, Mark Achtman, Zhemin Zhou

**Author notes:** These authors contributed equally (Ling Zhong, Heng Li, Shuting Xia, Shichang Xie, Raffaella Bianucci, Shengkai Li). Correspondence: Huilin Yang; Mark Achtman; Zhemin Zhou.

## Abstract

The human oral microbiome represents an intimate and enduring partnership between host and microbial communities, yet its evolutionary trajectory across deep time remains largely uncovered. Here we reconstruct the global history of the oral microbiome over 102,400 years using a global collection of 1854 oral samples spanning 61 countries. We reveal that major cultural transitions of the Neolithic Revolution, industrialization, and recent medical advances have driven a directional, globally coordinated co-evolution of oral microbial ecology, function, and population structure. Network analysis identifies two universal microbial modules of dental calculus modules: the commensal-enriched DCM1 module and the pathogen-enriched DCM2, both exhibiting progressive ecological polarization. DCM2 surged around 4,000 years ago with agricultural intensification but subsequently declined in modern European populations while persisting at high levels in African and some American groups, a geographic divide mirroring socioeconomic inequities. Species-level analyses demonstrate widespread directional changes, with obligate anaerobes declining as aerotolerant taxa expand, reflecting a carbohydrate-associated functional remodeling toward starch metabolism. Phylogenomic reconstruction reveals contrasting evolutionary trajectories. *Pauljensenia mediterranea* experienced near-extinction following a severe bottleneck, while *Actinomyces israelii* underwent a selective sweep with rapid population expansion 1,000 years ago. Critically, the oral resistome has accelerated in the antibiotic era, resulting in ∼33–50-fold surge of highly mobile, clinically relevant antimicrobial resistance genes in the last century, with *Streptococcus* acting as a central hub of dissemination. Collectively, these results position the oral microbiome as an active participant that has co-evolved with human subsistence, urbanization and medical practice, with direct implications for global oral health, antimicrobial resistance and interventions aimed at restoring resilient microbial states.

## Introduction

The human body hosts diverse microbial communities that influence physiology, immunity, and metabolism, yet their long-term evolutionary trajectories remain underexplored (*1*, *2*). Decoding host-microbe interactions is critical not only for reconstructing our biological past but also for understanding how they have influenced nutrition, immune function, and disease susceptibility across deep time (*3*).

The oral cavity serves as one of the most accessible and diverse environments for investigating long-term human–microbe interactions (*4*). Modern oral dysbiosis, characterized by a shift from commensal-to pathogen-enriched communities, are associated with various conditions from caries and periodontitis to systemic inflammatory diseases (*5*, *6*). Such disorders have been linked to “diseases of civilization”, increasing in prevalence with the developments of agriculture and industrialization (*7*, *8*). Yet the tempo, scale, and microbial mechanisms underlying these transitions remain obscure.

Recent advances in ancient DNA have revolutionized our capacity to reconstruct past microbial communities, with fossilized dental calculus preserving a durable archive of oral microbial DNA for 1000’s of years (*9*, *10*). Pioneering efforts revealed microbial responses to dietary shifts subsistence, and health status (*3*, *10*). However, existing studies have been constrained to discrete sites or historical periods, preclude detection of global evolutionary patterns and assessment of whether observed trends represent universal human adaptations or region-specific responses to localized pressures.

Understanding the evolutionary ecology of the oral microbiome is particularly important given its dual role as both a commensal ecosystem and a reservoir for pathogens and antimicrobial resistance genes (ARGs) (*11*, *12*). The oral cavity functions as both a barrier and a bridge, linking environmental exposures, social transmission, and systemic infection pathways and responding rapidly to diet, hygiene, and medical interventions (*13*). Yet whether these selective pressures have driven directional, predictable evolutionary trends across millennia, or merely transient fluctuations, remains unresolved.

Here we reconstruct the global evolutionary history of the human oral microbiome over the past 102,400 years, spanning 61 countries. Integrating 272 our ancient dental calculus samples with 1588 modern and historical datasets, this unprecedented temporal and geographic scope enable us to chart the global ecological, functional, and resistome dynamics of the oral microbiome. We identify long-persisting microbial modules that polarize into commensal– and pathogen-enriched states, reveal deep temporal and geographic structuring linked to agricultural and industrial transitions, and document a dramatic, recent acceleration of antimicrobial resistance. Together, these findings demonstrate that the oral microbiome has not been a passive passenger in human history but an active participant, with its evolution mirroring and mediating the transformations of human society itself.

## Results

### A 100,000-year global collection of the oral microbiome

We assembled a comprehensive global dataset comprising 1,854 oral microbiome samples from modern humans and Neanderthals spanning 102,400 years across 61 countries (**Figure 1a**). The dataset includes 272 our own ancient dental calculus samples from targeted excavations across Italy (n=77; 390–55,000 years BP), the United Kingdom (n=120; 208–2400 years BP), other European countries (n=5; 821–1371 years BP), and pre-Columbian Mexico (n=14; ∼1,000 years BP). Additionally, we sampled six and 46 modern dental plaque samples from China and Thailand, respectively. Integration with previously published datasets yielded substantial global representation, the final collection achieves global representation, encompassing samples from Americas (n=928), Europe (n=602), and Asia (n=268) **(Supplementary Table 1 and 2)**.

**Figure 1.**
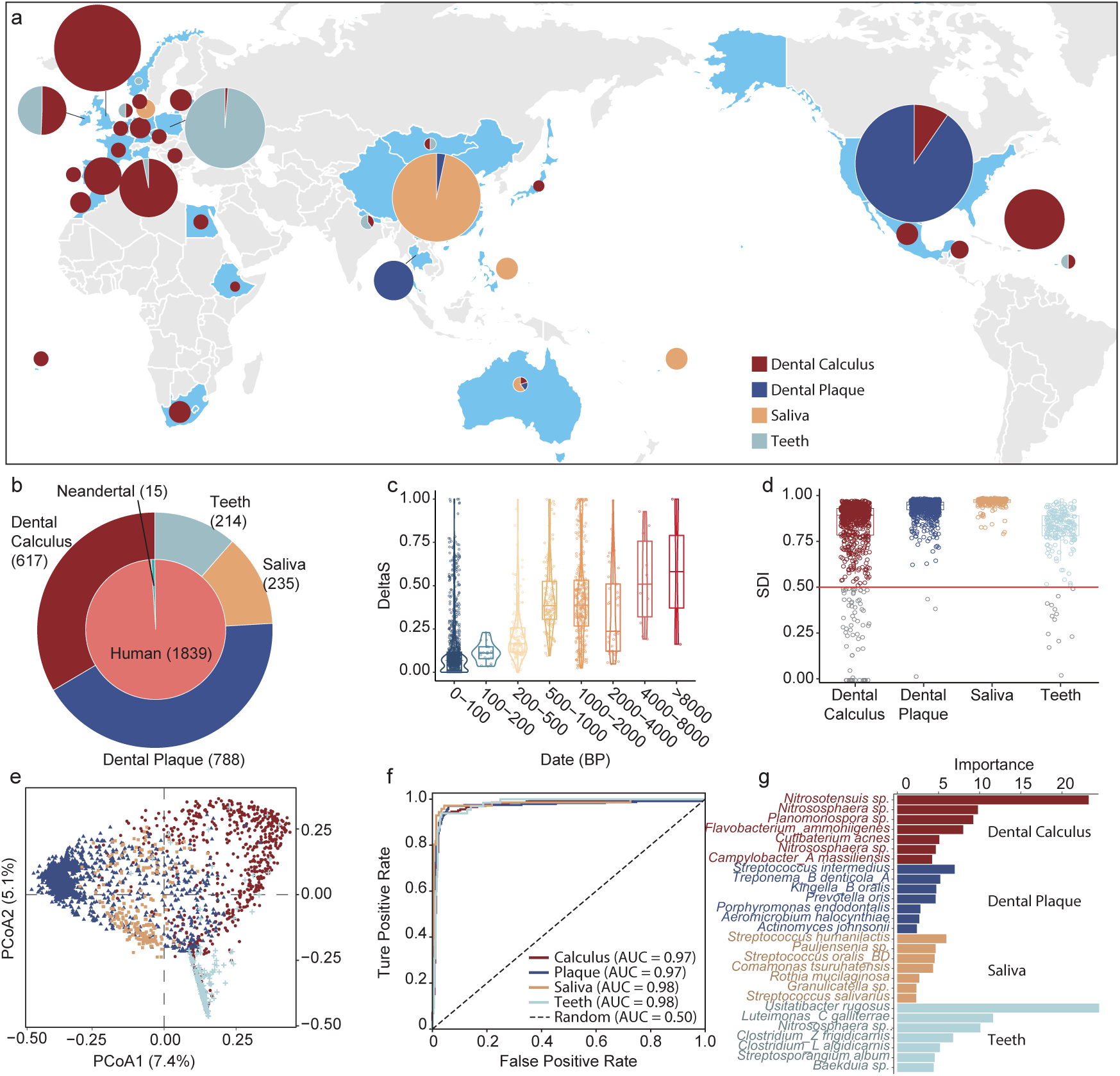
| Global oral microbiome dataset and quality control across anatomical sites and time. (**a**) Geographic and temporal distribution of 1,857 oral microbiome samples spanning 102,400 years across 61 countries. Pie charts indicate sample composition by type at each location, with circle size proportional to sample number (see scale). (**b**) Overall dataset composition showing distribution of sample types: dental plaque (n = 788), dental calculus (n = 622), saliva (n = 235), teeth (n = 212), human host (n = 1,846), and Neanderthal (n = 15). (**c**) Ancient DNA authentication via mapDamage analysis showing cytosine-to-thymine (C-to-T) deamination frequencies at 5′ fragment termini plotted against sample age. Violin plots demonstrate age-dependent increases in damage patterns. (**d**) Simpson Diversity Index (SDI) distributions across sample types. Most samples show high diversity (SDI > 0.5), while low-diversity outliers (SDI < 0.5), predominantly in dental calculus, indicate degradation or contamination. (**e**) Principal coordinates analysis (PCoA) based on Bray–Curtis dissimilarity showing distinct clustering by anatomical site. Marginal histograms display density distributions along PCo1 (7.4% variance) and PCo2 (4.3% variance). (**f**) Receiver operating characteristic (ROC) curves for multi-class logistic regression models predicting anatomical origin from microbial composition. Area under the curve (AUC) values of 0.97–0.98 demonstrate high classification accuracy across all sample types. (**g**) Feature importance analysis showing the top taxonomic contributors to anatomical site classification. Colors correspond to sample types in **b** and **e**.

Sample types comprised primarily dental plaque (n = 788) and dental calculus (n = 617), with additional contributions from saliva (n = 235) and dental tissues 图b 是 teeth (n = 214) (**Figure 1b**). To validate ancient DNA authenticity, we used mapDamage to assess cytosine-to-thymine substitution frequencies at 5′ fragment termini, a characteristic signature of post-mortem cytosine deamination (**Supplementary Figure 1**). The non-UDG-treated ancient samples exhibited age-correlated damage patterns, with samples exceeding 4,000 years showing the highest deamination levels (>50%) and most of modern samples (<200 years BP) displayed minimal damage (<2%), confirming the authenticity and suitability of the ancient DNA for evolutionary analyses (**Figure 1c**).

### Anatomical niche determines taxonomic composition and reveals contamination

Taxonomic profiling revealed diverse microbial taxa across oral sources. Saliva exhibited the greatest species richness per sample, while dental calculus contained the largest cumulative richness across all samples (**Supplementary Figure 2b/c**). Twenty most abundant species comprising 36.3% of total microbial abundance (**Supplementary Figure 2a**), comprising canonical oral colonizers like *Streptococcus* spp., *Actinomyces* spp., as well as opportunistic pathogens like *Tannerella forsythia* and *Haemophilus parainfluenzae*. Principal coordinates analysis (PCoA) demonstrated strong clustering by anatomical site, explaining 18.0% of overall variations (PEMANOVA test, **Figure 1e**). Samples from each anatomical niche form distinct clusters, prompting them as the primary determinants of microbial community structure.

Most oral microbiome samples harbored rich and evenly distributed communities, achieving Simpson Diversity Index (SDI) of >0.5 (**Figure 1d**). A minority of samples, primarily derived from ancient dentin, exhibited substantially lower diversity (SDI < 0.5), reflecting extensive DNA degradation or environmental contamination. These low-diversity samples were excluded from downstream analyses to ensure data integrity.

To objectively evaluate sample reliability and detect contamination, we applied logistic regression models to predict anatomical origin based on microbial composition. The models achieved high accuracy (0.96 – 0.99) and AUC (0.97 – 0.98) across all four sites (**Figure 1f**). Key taxonomic signatures effectively distinguished the sample types. Notably, ammonia-oxidizing archaea such as *Nitrososphaera* and *Nitrosotenuis* were enriched in dental calculus, whereas a diversity of *Streptococcus* species were characterized in plaque and saliva (**Figure 1g**). A slight misclassification of dentin as calculus (4%) was observed, possibly indicating post-mortem infiltration of calculus-associated taxa. We excluded samples with low classification confidence (prediction probability <0.6), yielding a final high-quality dataset of 1735 samples (**Supplementary Table 2**).

### Dental calculus reveals distinct temporal and geographic structure

Focusing on dental calculus, the most reliably preserved oral substrate, we analyzed 606 samples spanning the full temporal range. Higher SDI were observed in European samples and within the last 200 years compared to others (**Figure 2a–b**). Uniform Manifold Approximation and Projection (UMAP) further confirmed distinct clustering by temporal and geographic origin (**Figure 2c–d**), underscoring their roles in shaping oral microbiome composition.

**Figure 2.**
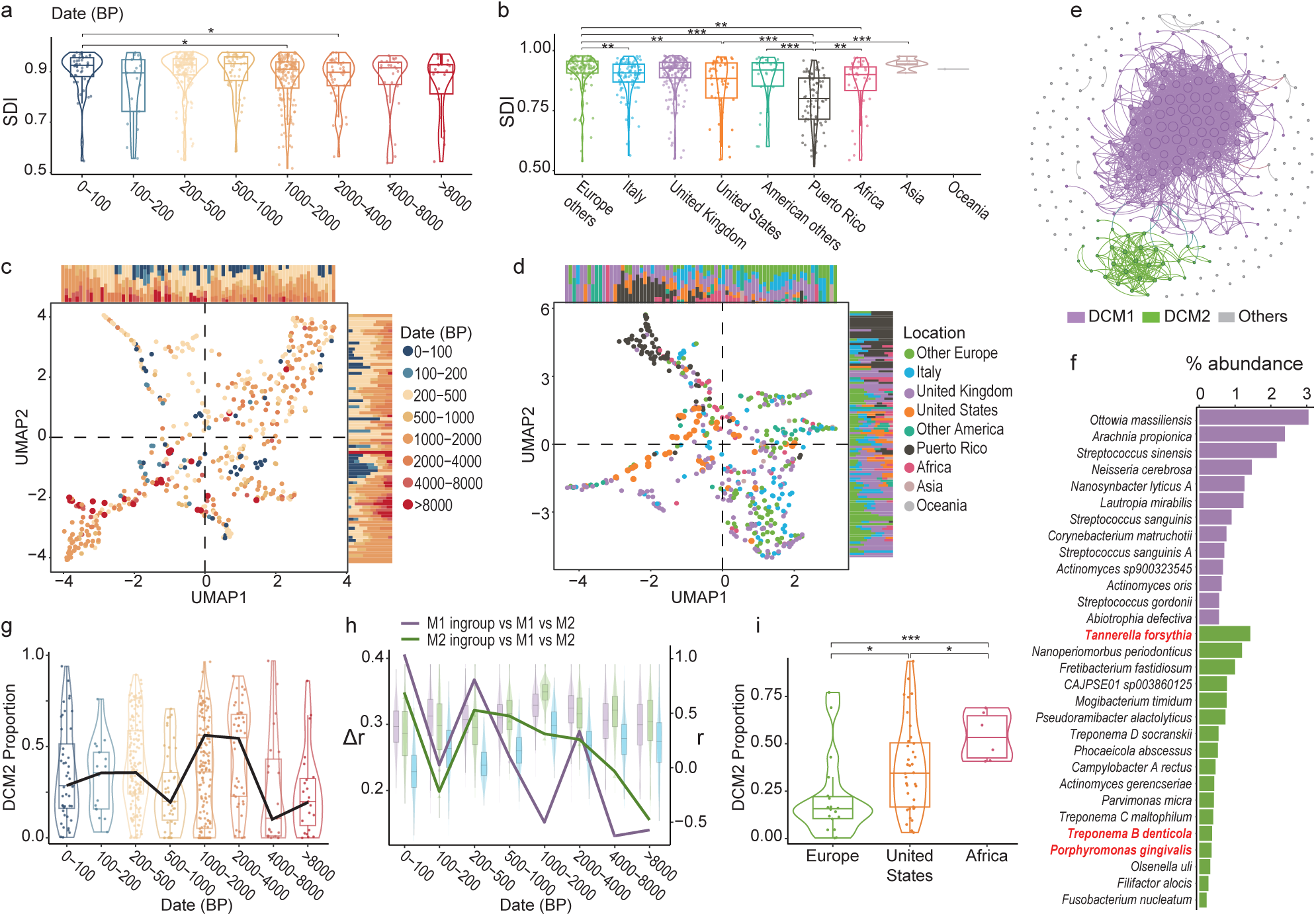
| Spatiotemporal structure of the dental calculus microbiome reveals modular reorganization and health inequities. (**a**) Temporal variation in Simpson Diversity Index (SDI) across eight time periods. Asterisks indicate significant differences between groups (two-sided Wilcoxon tests with Bonferroni correction). **(b)** Geographic variation in SDI across nine regions. (**c**) Uniform Manifold Approximation and Projection (UMAP) of dental calculus samples colored by sample age (years BP). Marginal bar plots show temporal distributions along each axis. (**d**) UMAP colored by geographic origin demonstrates spatial clustering. Marginal bar plots display regional distributions. (**e**) Co-occurrence network analysis identifies two distinct dental calculus modules (DCMs): DCM1 (purple) comprises predominantly commensal species, while DCM2 (green) is enriched for periodontal pathogens. Edge width represents correlation strength; node size reflects taxon prevalence. (**f**) Taxonomic composition of DCM1 and DCM2. DCM1 contains health-associated commensals (*Prevotella*, *Capnocytophaga* species), whereas DCM2 harbors disease-associated taxa including the “red complex” pathogens (*Porphyromonas gingivalis*, *Tannerella forsythia*, *Treponema denticola*, shown in red). (**g**) Historical trajectory of DCM2 prevalence over 102,400 years. DCM2 abundance increased sharply around 4,000 BP, peaking at >50% between 2,000–4,000 BP, before declining to ∼30% in modern samples. Box plots show median, quartiles, and range; violin plots show probability density. (**h**) Temporal dynamics of inter– and intra-module correlations. Bar plot shows median within-module correlations for DCM1 (purple) and DCM2 (green) versus between-module correlations (blue) across time periods. Violin plots display correlation value distributions for each comparison type. The line graph (right y-axis) shows the difference in average correlation values (within minus between modules), revealing progressive decoupling from Δ = 0.14 at ∼8,000 BP to Δ = 0.51 in modern samples. (**i**) Contemporary geographic distribution of DCM2 prevalence. African populations show significantly elevated pathogen-module representation (∼50%) compared to European cohorts (∼10%) (two-sided Wilcoxon tests). Statistical significance: \**p* < 0.05, \*\**p* < 0.01, \*\*\**p* < 0.001.

We employed random forest models to predict the spatiotemporal origin of samples based on their taxonomic profiles, which achieved robust performance in predicting temporal periods (AUC = 0.90) and geographic regions (AUC = 0.89), respectively (**Supplementary Figure 3a-c**). A total of 14 taxa significantly contributed to the temporal classification, while 12 contributed geographically. Notably, eight of these informative taxa overlapped, underscoring the tight correlation between temporal and geographic factors (**Supplementary Figure 3d-f**).

We investigated regional connectivity using the network analysis of sample-to-sample Spearman’s correlations (**Supplementary Figure 4a**). Puerto Rican samples formed an isolated cluster, likely reflecting geographic isolation and unique environmental conditions. In contrast, European and Asian samples exhibited high connectivity, evidenced by their substantially greater eigen centralities (European: 0.23, Asia: 0.25) and average inter-regional connections (European: 10.8, Asia: 8.8) (**Supplementary Figure 4b-c**). This high Eurasian connectivity likely reflects sustained population mobility, trade networks, and cultural exchange throughout history, which maintained relatively homogeneous oral microbiomes across national boundaries.

### Millennial-scale ecological polarization of dental calculus microbial modules

Network analysis of species co-occurrence patterns in dental calculus samples identified two robust modules, which we designated as dental calculus modules (DCMs), that universally present in almost all temporal and spatial subgroups (**Figure 2e, Supplementary Figures 5, and Supplementary Table 3**). DCM1 comprised predominantly commensal taxa linked to oral health including *Prevotella* and *Capnocytophaga* species (**Figure 2f**). In contrast, DCM2 was enriched with potential periodontal pathogens, most notably the three “red complex” bacteria (*Porphyromonas gingivalis, T. forsythia,* and *Treponema_B denticola*), as well as numerous additional disease-associated taxa.

Temporal analysis revealed substantial shifts in module prevalence (**Figure 2g**). The pathogen-associated DCM2 rose sharply around 4000 BP, from a median of 25.1% in earlier periods to over 45.7% between 1000 and 4000 BP, before gradually declining to 34.4% in the present day. This trend was further confirmed in a European sub-analysis, where DCM2 prevalence dropped to 23.4% in the last century (**Supplementary Figure 6a**).

Critically, inter-module correlation analyses demonstrated the progressive functional and ecological decoupling of DCM1 and DCM2 over time, both globally and in Europe (**Figure 2h, Supplementary Figure 6b**). Around 8000 BP, taxa between modules showed a weak positive correlation (*r* = 0.18 vs. 0.32 intra-group, Δr = 0.14), which steadily weakened and transitioned to a strong negative correlation in modern samples (r = –0.16 vs. 0.35, Δr = 0.51). The most pronounced decline in correlation coincided with the peak in DCM2 prevalence around 4000 BP (r = 0.26 vs. 0.55, Δr = 0.29), indicating that the oral microbiome has undergone a long-term ecological polarization, separating into distinct commensal and pathogen-enriched states.

Analysis of contemporary populations revealed global inequality in terms of the DCM1–DCM2 balance **(Figure 2i)**. The commensal-dominated DCM1 communities are more common in European and certain American cohorts (21.5%), whereas the pathogen-enriched DCM2 remains prevalent at high levels (55.4%) in populations across Africa and parts of the North Americas. This stark geographical divide reveals that dysbiotic oral ecosystems have disproportionately persisted in vulnerable populations. Such patterns mirror underlying socioeconomic disparities, suggesting that health inequities may have left a lasting biological imprint on the human microbiome.

### Species-level temporal trends reveal widespread directional change with geographic bias

To quantify individual species’ contributions to long-term temporal shifts, we modeled associations between microbial relative abundances and sample age using generalized linear models, adjusting for geographic origin (**Figure 3a**). This analysis identified significant temporal dynamics in 144 of 310 tested species (46.8%), indicating widespread temporal shifts in taxonomic composition of oral microbiome. Subsequent classification with Gaussian Mixture Models delineated 63 taxa as increasing and 81 as decreasing in abundance over time (**Figure 3a/b, Supplementary Figure 7a, and Supplementary Table 4**).

**Figure 3.**
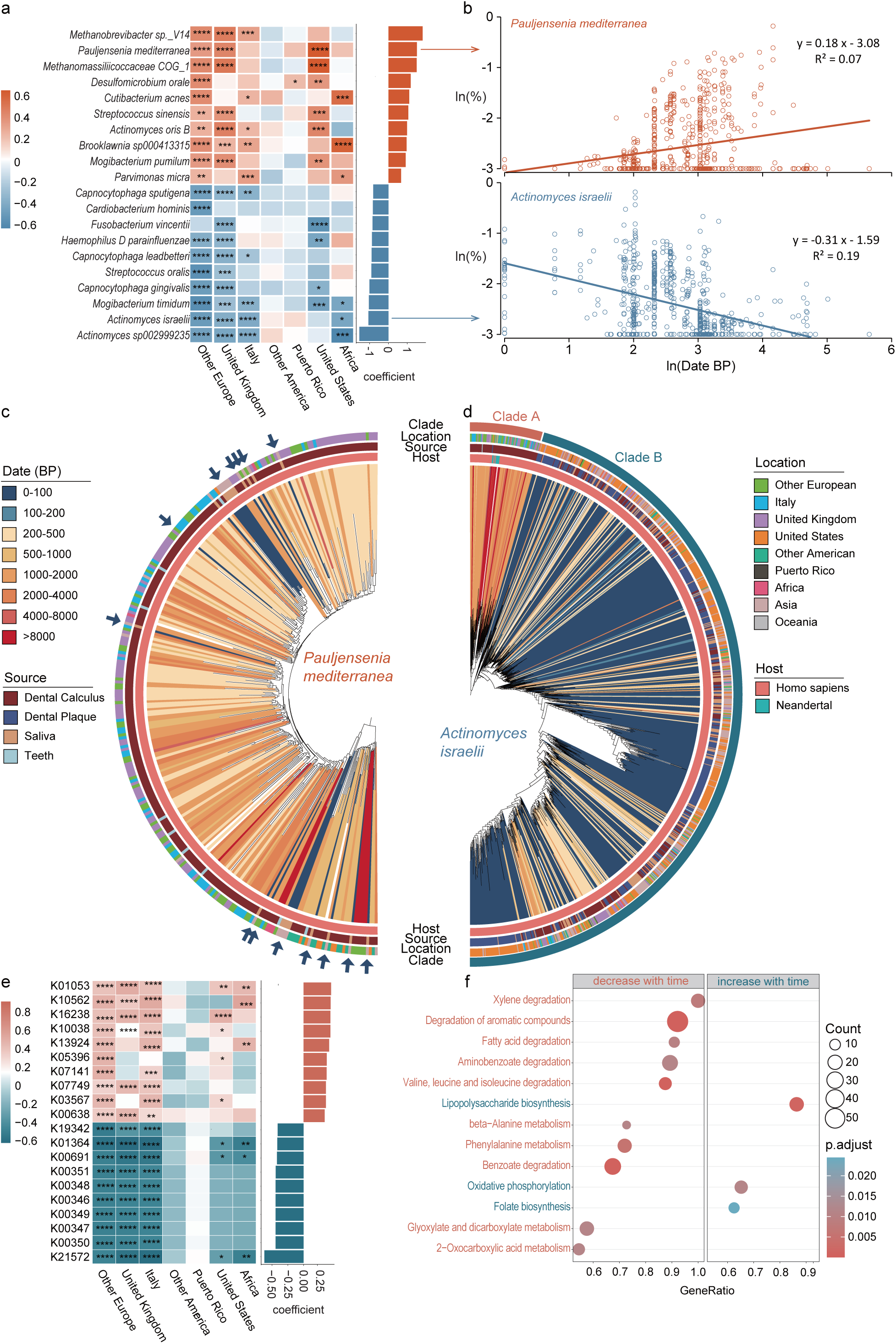
| Species-level temporal dynamics reveal directional shifts, geographic heterogeneity, and functional adaptation. (**a**) Heatmap showing coefficients from generalized linear models testing associations between species abundance and sample age across geographic regions. Each row represents one species with significant temporal trends; columns represent geographic regions (Europe, United Kingdom, Italy, North America, Puerto Rico, United States, Africa). Color intensity indicates coefficient magnitude (red = positive/increasing, blue = negative/decreasing); asterisks denote statistical significance (\**q* < 0.05, \*\**q* < 0.01, \*\*\**q* < 0.001, \*\*\*\**q* < 0.0001, two-sided Wald tests with FDR correction). (**b**) Trajectory plot showing temporal trends for two representative species over log[[-transformed sample age. *Pauljensenia mediterranea* (coral, top) exhibits stable or slightly declining abundance, while *Actinomyces israelii* (blue, bottom) shows consistent increase over time. (**c-d**) Phylogenetic trees with metadata rings for *P. mediterranea* (c) and *A. israelii* (d). Tree tips represent individual strains colored by sample age (innermost ring, see legend). Outer rings indicate geographic location, sample source (dental calculus, plaque, saliva, teeth), and host species (human vs. Neanderthal). *P. mediterranea* shows broad phylogenetic diversity across ancient samples with recent population contraction, with the contemporary samples pointed by blue arrows. *A. israelii* exhibits a distinct clade B expansion in modern samples (dark blue cluster, <200 BP), indicating a selective sweep beginning ∼1,000 BP. Branch lengths represent genetic distance; metadata rings provide spatiotemporal context for evolutionary patterns. (**e**) Heatmap showing temporal coefficients for 21 functional genes across the same geographic regions, demonstrating pervasive but geographically heterogeneous temporal trends. KEGG orthology (KO) identifiers indicate functional genes. (**f**) KEGG pathway enrichment analysis comparing ancient versus modern samples. Dot plot shows gene ratio (x-axis) and adjusted P-values (color) for significantly enriched pathways. Dot size represents gene count. Pathways enriched in ancient samples (left, <0.9 GeneRatio) include aromatic compound degradation, fatty acid metabolism, and amino acid degradation. Modern-enriched pathways (right, >1.0 GeneRatio) include carbohydrate metabolism and oxidative phosphorylation.

We observed notable increases in the relative abundance of *Streptococcus* species and *Actinomyces israelii*, alongside consistent declines in obligate anaerobes such as *Methanobrevibacter* and *Capnocytophaga* species. This pattern suggests a long-term ecological transition toward more aerotolerant oral microbiome. However, the temporal trends varied in different geographic regions. Regional stratification analysis showed that European populations exhibited more pronounced and consistent changes compared to African and American populations (**Figure 3a and Supplementary Table 4**).

### Phylogenetic analysis reveals population structure changes and expansion events

We reconstructed phylogeny for two species with the most significant temporal trends, illustrating their contrasting evolutionary trajectories associated with cultural transitions. *Pauljensenia mediterranea*, once widespread across Eurasia and the Americas, experienced a severe population bottleneck over the past 200 years (**Figure 3b/c**). Compared to its broad phylogenetic diversity in ancient samples and widespread presence across geographic regions, the species was now primarily only observed in saliva samples in Asian population, leaving genetically restricted clades that accounted for only 7.9% of the overall diversity across the phylogeny (**Supplementary Figure 8a, b**).

In contrast, *A. israelii* experienced a selective sweep starting around 1,000 years ago, accompanied by its rapid population expansion. The species’ phylogeny consisted of two major clades (A and B) diverging directly from the root (**Figure 3d**). Clade A consisted of 84 ancient samples originating between 208 and 102,400 BPs, including 7 of the 15 Neanderthals (**Supplementary Figure 8c**), demonstrating its historical universal presence across Eurasia. This clade, however, experienced rapid decrease in its relatively frequencies after 1000 BP and was not observed in the past 200 years. Instead, Clade B, only occasionally found in 2000 BP, emerged with time and became the predominant clade in the past 1000 years. This clade replacement event coincided with intensified agriculture and urbanization, suggesting ecological adaptation to altered human lifestyles (**Supplementary Figure 8d**).

### Carbohydrate-associated functional remodeling of the oral microbiome

We applied temporal analysis to KEGG pathway annotations, which identified 1343 and 965 KOs that down-or up-regulated with time, indicating profound metabolic rewiring of the dental calculus microbiome (**Supplementary Table 5**). Specifically, starch-binding outer membrane protein gene K21572 exhibited the most dramatic upregulation in modern samples across all geographic regions (**Figure 3e)**, providing evidence for increased dietary starch intake following agricultural transitions.

Conversely, genes involved in anaerobic metabolism exhibited consistent declines (**Figure 3f**). Pathways for amino acid metabolism and aromatic compound degradation were enriched in ancient samples, possibly reflecting pre-agricultural, meat-rich diets. In contrast, modern samples showed enrichment for oxidative phosphorylation pathways and increased lipopolysaccharide synthesis, indicating a shift toward aerobic energy metabolism in contemporary oral microbiomes.

### Geographic and temporal breakpoint analysis reveals cultural transition signatures

Breakpoint analysis of species abundance trajectories revealed pivotal periods of accelerated oral ecological evolution (**Figure 4a-b, Supplementary Figure 9**). Initial analysis of European samples (n=247) identified two periods with excessively high level of breakpoints at approximately 5000 BP and 200 BP (**Figure 4c**), These breakpoint clusters temporally coincident with the European Neolithic and Industrial Revolutions, respectively, indicating close associations between major cultural transitions and reconstructions of oral microbial communities.

**Figure 4.**
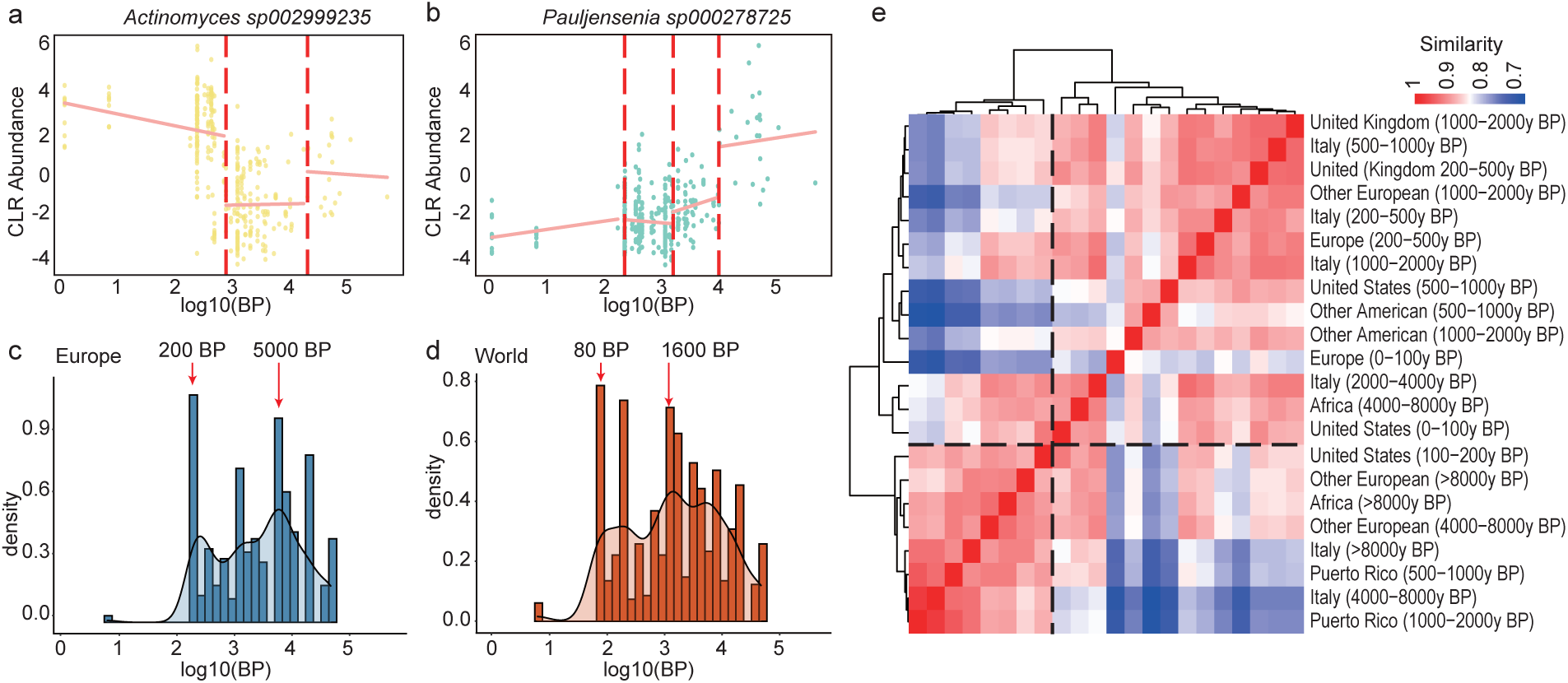
| Breakpoint analysis identifies cultural transition signatures in oral microbiome evolution. (**a**–**b)** Representative examples of piecewise regression models detecting temporal breakpoints in species abundance trajectories. Yellow points represent individual samples; the fitted piecewise linear model (coral line) captures distinct phases of decline and stabilization. **(c**) Temporal distribution of breakpoints identified across species using European samples only (n = 247). Histogram bars show the density of breakpoints across log[[-transformed age bins; the overlaid kernel density curve (black line) reveals two prominent peaks at ∼4,000 BP and ∼200 BP, corresponding to the Neolithic Revolution and Industrial Revolution, respectively. (**d**) Global breakpoint distribution using the full dataset with geographic adjustment. Histogram (coral bars) and density curve (black line) show breakpoint clustering at ∼200 BP and ∼2,000 BP. The ∼200 BP peak reflects the globally synchronous impact of industrialization, while the broader ∼2,000 BP signal suggests temporally heterogeneous agricultural transitions across continents, with delayed Neolithic adoption in the Americas compared to Eurasia. Density values (y-axis) represent the proportion of breakpoints occurring within each time bin. (**e**) Hierarchical clustering heatmap of cross-regional-temporal similarity based on centroid community structures. Rows and columns represent 27 geographic-temporal population groups (format: Location (AgeRange)). Color intensity indicates Bray–Curtis similarity (red = high similarity, blue = low dissimilarity; log[[-transformed scale shown at top right). Dendrogram reveals a primary temporal bifurcation separating ancient samples (>4,000 BP, upper-left cluster including Africa_8000+, Europe_8000+, and Puerto Rico samples) from recent/modern populations (lower-right cluster).

Subsequent global analysis, adjusted for geographic confounding factors, confirmed these evolutionary pulses while revealing distinct regional chronologies. Global breakpoints clustered around 1600 BP and 80 BP, reflecting the delayed Neolithic and Industrial transition influences, particularly in the Americas where most samples were from (**Figure 4d**). This temporal disparity aligns with the independent and later establishment of agricultural systems in the New World, demonstrating how major cultural transitions reshaped oral microbiomes along distinct spatiotemporal trajectories.

### Hierarchical clustering reveals deep temporal and geographic structuring of the oral microbiome

To further elucidate the spatial and temporal drivers of microbial community variation, we performed hierarchical clustering on centroid community structures of geographically and temporally distinct populations. This analysis revealed a deep temporal bifurcation, separating a clade of ancient samples (primarily > 4,000 BP) from a clade encompassing more recent and modern populations (**Figure 4e**). This primary temporal split, which transcends geographic boundaries, is strikingly evident in the tight clustering of the pre-neolithic African and European samples (e.g., Africa_8000+y and Europe_8000+y), suggesting a conserved ancestral microbiome state that predates major dietary and lifestyle transitions.

Within the more recent clade, strong secondary clustering emerged along regional and historical lines. For example, Italian samples spanning 500–2000 BPs grouped together, revealing remarkable temporal continuity within a single geographic context despite cultural transitions. Similarly, very recent North American and European samples (0–100y) form a distinct subcluster, separated from older European (200–500y) and pre-colonial North American populations. These patterns suggest that while the initial divergence in community structure was driven by ancient, globally transformative events, subsequent layering of selective pressures, including regional dietary traditions, colonial admixture, and industrial-era hygiene and nutrition, has shaped the fine-scale diversification observed today.

### The ongoing reshaping of oral microbiome towards increased resistance in the antibiotic era

These long-term patterns raise the question of how oral microbiomes are continuing to evolve today, particularly under the selective pressures imposed by modern medicine. To address this, we profiled ARGs using the Structured ARG (SARG) database (*14*), which classifies genes into four ranks by mobility and clinical relevance, with Ranks I and II include highly mobile, clinically urgent ARGs frequently located on plasmids or transposons.

This framework revealed a dramatic temporal divergence in the oral resistome (**Supplementary Table 6)**. While the overall abundance of ARGs increased steadily over time, the most significant shift occurred in the last century (**Figure 5a**). The low-mobility, intrinsic resistome (Ranks III-IV) showed only modest, gradual increases (2.5–6.3-fold), whereas the highly mobile and clinically significant ARGs (Ranks I–II) escalated dramatically, surging 33–50-fold during the last 100 years. This unprecedented acceleration coincides with the antibiotic era, reflecting the transformative selective pressures of modern medicine and hygiene practices on the oral microbiome.

**Figure 5.**
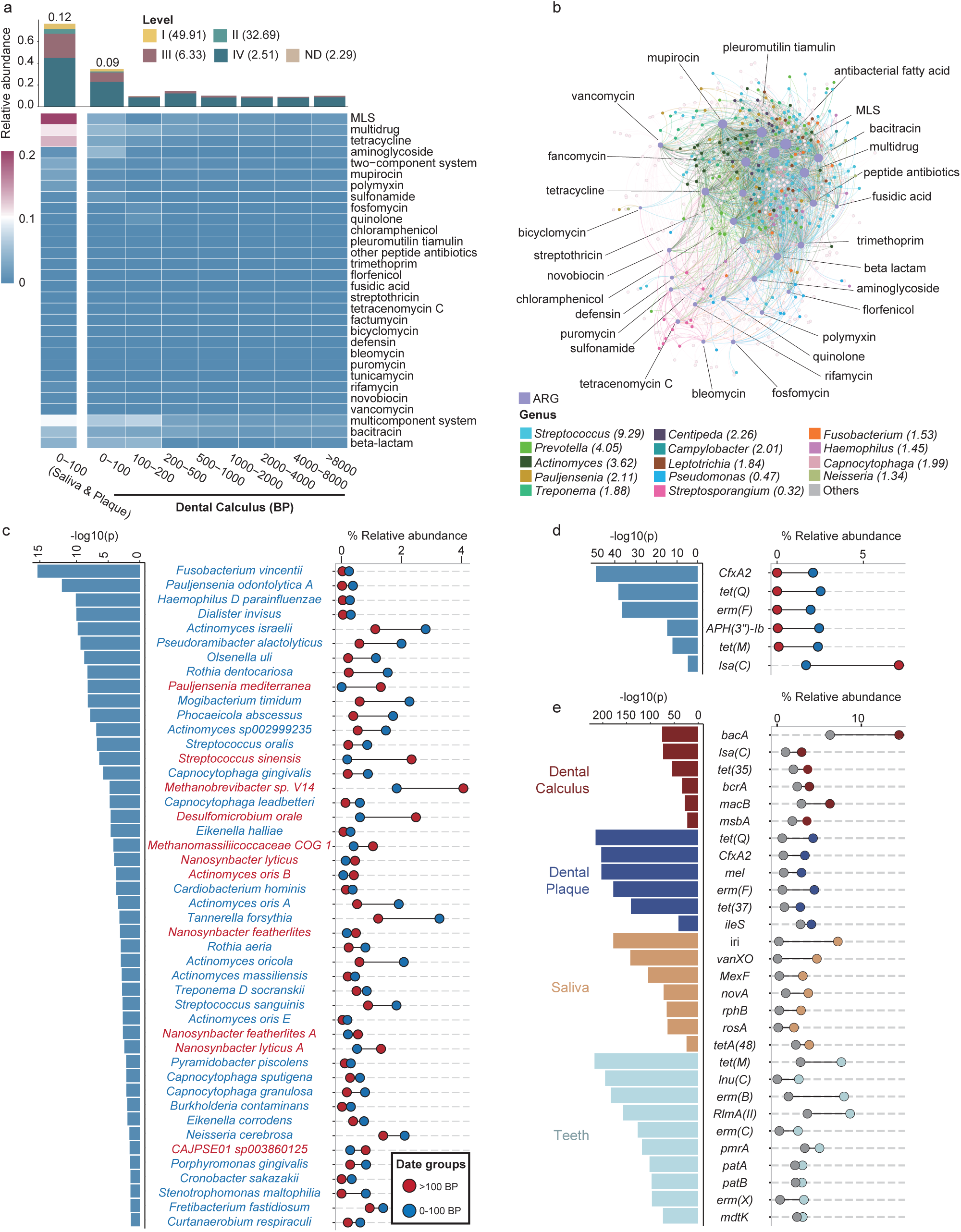
| Antimicrobial resistance gene expansion in the modern oral microbiome driven by mobile elements and *Streptococcus* enrichment. **(a)** Temporal dynamics of antimicrobial resistance gene (ARG) abundance across 102,400 years. Top stacked bar chart shows the proportion of ARG ranks by clinical mobility and urgency: Rank I (highly mobile, clinically urgent, n = 49,910 detections), Rank II (mobile, clinically relevant, n = 32,690), Rank III (chromosomal, moderate mobility, n = 6,330), Rank IV (intrinsic, low mobility, n = 2,510), and unassessed genes (n = 1,560). Heatmap displays relative abundance of 33 specific ARG types (rows) across eight temporal bins (columns), with macrolide–lincosamide–streptogramin (MLS), tetracycline, and β-lactam resistance genes showing pronounced modern enrichment. Leftmost bars show proportion breakdown by saliva and plaque samples in the 0–100 BP period. (**b**) Co-abundance network of ARG-harboring taxa. Nodes represent bacterial genera sized by total ARG count and colored by taxonomy. Legend shows the total eigencentrality scores of each genus. Edges connect genera with correlated ARG profiles; edge thickness reflects correlation strength. (**c**) Linear discriminant analysis (LDA) effect size (LEfSe) comparing species enrichment in ancient (>100 BP, blue bars) versus modern (<100 BP, coral bars) dental calculus samples. LDA scores >2 indicate significant differential abundance. (**d**) LEfSe analysis of ARG enrichment in modern samples by anatomical source. Bar length indicates LDA score; genes are grouped by antibiotic class (labels in parentheses). Saliva samples (tan bars) show strong enrichment for macrolide resistance genes including *erm*(B), *erm*(F), *erm*(C), and aminoglycoside resistance genes (*aph*, *ant*). (**e**) LEfSe showing dental calculus (dark coral), dental plaque (burgundy), saliva (tan), and teeth (light blue) source-specific ARG enrichment patterns. Dental plaque is particularly enriched for β-lactam resistance genes (*bac*, *bla*, *bcr*) and tetracycline resistance (*tet*(35), *msb*, *mac*).

Co-abundance network analysis revealed *Streptococcus* as the central hub of ARGs dissemination (eigenvector centrality score = 9.29), far surpassing *Prevotella* (4.05) and *Actinomyces* (3.62) (**Figure 5b**). *Streptococcus* species not only harbored a wide spectrum of resistance determinants but also maintained dense network connectivity, positioning them as critical reservoirs and facilitators of horizontal gene transfer. Importantly, multiple *Streptococcus* species show enrichment specifically in the last century together with other potential oral pathogens (red complex, *Eikenella corrodens*, etc.), suggesting that their expansion may underlie the sharp rise of ARGs prevalence (**Figure 5c**).

### High ARGs levels in modern plaque and saliva confirm evolutionary acceleration

Contemporary plaque and saliva samples exhibited the highest ARGs abundances in the entire dataset (**Figure 5a; Supplementary Table 6**), confirming the accelerating pace of resistome evolution. Saliva samples were particularly enriched for rank I macrolide resistance genes (*erm*(B), *erm*(C), *erm*(X)), while modern plaque samples showed elevated levels of tetracycline resistance genes (*tet*(Q)) (**Figure 5d/e**). This pattern underscores the oral cavity’s role as both a reservoir and transmission route for clinically relevant antimicrobial resistance.

Together, our findings demonstrate that the rate of ARGs accumulation has accelerated by orders of magnitude in the modern era, fundamentally altering the genetic architecture of the oral microbiome within a handful of generations. The co-enrichment of opportunistic pathogens, including multiple *Streptococcus* lineages, and ARGs illustrates a potential feedback loop between clinical interventions and resistome amplification, a broader “diseases of civilization” wherein industrial-era practices reshape microbial ecosystems toward pathogenic potential.

## Discussion

Our reconstruction of the human oral microbiome across 100,000 years provides unprecedented evidence that this microbial ecosystem serves as a high-fidelity archive of human cultural and technological evolution. By assembling the largest and most diverse spatiotemporal dataset of oral metagenomes to date, we demonstrate that major transitions in human history, from the adoption of agriculture to the Industrial Revolution and the antibiotic era, are indelibly recorded in the composition, function, and genomic evolution of our microbial companions. These findings reposition the oral microbiome from a passive passenger to a dynamic, co-evolving system that has been continuously reshaped by human behavior. These results move beyond prior small-scale studies of dental calculus to establish a global, temporally anchored framework, positioning the oral microbiome as a durable biosocial archive of human history.

A central discovery of our study is the long-term ecological polarization of the oral microbiome, characterized by the progressive decoupling of a commensal-dominated module (DCM1) and a pathogen-enriched module (DCM2). The sharp increase in DCM2 prevalence around 4,000 years BP in Europe coincides with intensified agriculture and urbanization, consistent with archaeological evidence of increased caries and periodontal disease during agricultural intensification (*15*). This temporal shift aligns with the “diseases of civilization” hypothesis proposing that Neolithic dietary shifts toward fermentable carbohydrates created selective pressures favoring acidogenic and aciduric pathogens (*16*).

However, our finding that inter-module correlations shifted from weakly positive (r = 0.18) to strongly negative (r = –0.16) over this period suggests a more complex process than simple pathogen enrichment. This ecological polarization may reflect the emergence of alternative stable states within oral biofilms, extends the bistability observed in other complex microbial ecosystems, where positive feedback loops reinforce divergent community configurations (*17*). The subsequent decline in DCM2 prevalence in modern European populations, contrasting with its persistence at ∼50% in contemporary African and certain American cohorts, possibly associated with the inequitably distributed health benefits of industrialization (*18*). This finding extends ancient microbiome research beyond taxonomic surveys (*3*) to establish that stable, module-level ecological organizations can persist, diversify, and polarize in parallel with human cultural evolution.

The dramatic population contraction of *P. mediterranea* and selective sweep of *A. israelii* over the past millennium exemplify how cultural transitions have reshaped species-level evolution. The replacement of historical *A. israelii* lineages (Clade A) with a derived clade (Clade B) after agricultural intensification suggests adaptive evolution to altered nutritional landscapes. *Actinomyces* species feature advanced starch-binding and degradation systems, mirroring our observed upregulation of starch-binding outer membrane protein genes (K21572) and supporting carbohydrate-induced functional changes (*3*). This aligns with experimental evidence of rapid microbiome responses to dietary shifts (*19*), extending such dynamics over evolutionary periods. Conversely, the near-extinction of *P. mediterranea*, an obligate anaerobe previously widespread across continents, may reflect the broader ecological shift toward aerotolerant communities that we observe in declining *Methanobrevibacter* and *Capnocytophaga* abundances. This likely stems from heightened oxygen exposure via refined diets, sugar-induced salivary flow, and hygiene practices eroding anaerobic habitats. Similar aerobic shifts have been documented in gut microbiomes of industrialized populations (*20*), implying shared pressures across body sites.

By identifying distinct temporal breakpoints in microbial community turnover, our study moves beyond correlation to establish a robust timeline linking microbial evolution to specific cultural transitions. The clustering of oral microbiome shifts around ∼4,000 BP and ∼200 BP mirrors the Neolithic and Industrial Revolutions, aligning microbial ecology with milestones in subsistence and technology. These results resonate with genomic evidence that large-scale cultural transformations leave parallel signatures in both host genomes and commensal microbiota (*21*).

Importantly, the temporal lags we identify between Europe and the Americas reinforce the global heterogeneity of these processes, emphasizing that while selective pressures were universal, their timing was mediated by regional histories of agriculture and industrialization, their impact on the microbiome followed the staggered timeline of human cultural diffusion, a pattern previously inferred from archaeological and human genetic data but never before observed at the microbial level (*22*).

The persistence of regional signatures despite global cultural convergence underscores the dual imprint of universal and local forces in microbial evolution. While industrialization produced synchronized ARG expansions across continents, oral microbiomes in geographically isolated populations, such as Puerto Rico, retained distinctive ecological structures. This pattern parallels observations in gut microbiomes, where industrialization drives broad functional convergence yet geography continues to explain taxonomic differentiation (*23*). Our results suggest that oral microbial evolution reflects a balance between homogenizing cultural forces and regionally specific ecological constraints.

The most striking signal of recent microbial evolution is the explosive proliferation of mobile antimicrobial resistance genes (ARGs) within the past century. The 33-50-fold increase in rank I–II ARGs dwarfs the scale of all prior evolutionary change observed in this study, underscoring the unprecedented selective pressure exerted by modern medicine. The centrality of *Streptococcus* in ARG dissemination and the highest ARG abundances in contemporary plaque and saliva highlights the oral cavity as both a reservoir and hub for horizontal gene transfer (*24*), consistent with clinical evidence of oral streptococci as major contributors to the resistome, posing a direct threat to public health (*12*). The co-enrichment of multiple *Streptococcus* lineages alongside ARGs in modern samples suggests a potential feedback loop wherein antibiotic-driven selection simultaneously favors resistant strains and pathogens capable of colonizing dysbiotic environments. This finding serves as a stark molecular signature of the Anthropocene, demonstrating how profound technological innovation can reshape microbial ecosystems on a global scale within just a few generations.

Our study also highlights critical limitations and avenues for future work. Although ancient DNA authenticity was rigorously validated, biases in preservation, geographic sampling, and chronological density inevitably shape the available record. In particular, African and Asian ancient datasets remain underrepresented, limiting resolution into non-European microbiome histories. Additionally, while functional inferences from metagenomes are robust, direct experimental validation of microbial metabolic shifts across deep time remains infeasible. Integrating stable isotope data, host genomic evidence, and functional assays of resurrected ancient enzymes may help bridge these gaps in the future (*25*).

Together, our 100,000-year reconstruction reveals that the human oral microbiome has undergone directional, globally coordinated evolutionary changes in response to agricultural and industrial transitions, with the antibiotic era triggering unprecedented resistome expansion. These findings establish the oral microbiome as an active participant in human biocultural evolution, whose trajectory reflects and mediates the biological consequences of societal transformation. As humanity continues to reshape microbial selection pressures through dietary choices, antibiotic use, and environmental modifications, understanding these long-term evolutionary patterns becomes essential for predicting future microbiome trajectories and developing interventions that promote microbial community resilience rather than further dysbiosis.

## Methods

### Sample collection and dataset assembly

The global oral microbiome dataset was compiled from newly generated and previously published data, comprising a total of 1,854 samples from modern and ancient humans and Neanderthals, spanning 102,400 years and 61 countries (details in **Supplementary Table 1**). This study includes 272 newly sequenced ancient dental calculus samples from four archaeological contexts: Italy (n=77; 390–55,000 years BP), the United Kingdom (n=120; 208–2,400 years BP), other European sites (n=5; 821–1,371 years BP), and pre-Columbian Mexico (n=14; ∼1,000 years BP). These were combined with 1,588 previously published oral microbiome samples obtained from public repositories including the NCBI Sequence Read Archive and European Nucleotide Archive, with accession numbers listed in Supplementary Table 1. Temporal categorization was performed as follows: Ancient (>8,000 years BP), Neolithic (4,000–8,000 years BP), Transitional (2,000–4,000 years BP), Historical (200–2,000 years BP), Recent (100–200 years BP), and Modern (<100 years BP). Modern sample data were obtained from publicly available databases with appropriate attribution to original studies as detailed in **Supplementary Table 1**.

### Ancient DNA extraction, library preparation, and sequencing

All ancient sample processing was conducted in a dedicated ancient DNA (aDNA) clean room facility at the University of Warwick under positive pressure with HEPA filtration, following established contamination prevention protocols. Dental calculus samples were mechanically cleaned to remove surface sediments using sterile dental tools, then decontaminated by UV irradiation (10 J/cm²) and a brief immersion in 0.5% sodium hypochlorite, followed by rinsing with DNA-free water. Samples were powdered using a sterile mortar and pestle. DNA was extracted from 10–50 mg of calculus powder using a silica-based column protocol optimized for recovering short, fragmented DNA (*26*). Negative extraction controls (no template) were processed in parallel with every batch of samples to monitor for laboratory contamination.

Double-stranded, non-UDG (uracil-DNA glycosylase) treated sequencing libraries were prepared from the extracted DNA to preserve aDNA damage patterns for authenticity assessment, following the protocol of Meyer and Kircher (*27*). Libraries were amplified with indexed primers, pooled, and subjected to shotgun sequencing on an Illumina NovaSeq 6000 platform, generating 150 bp paired-end reads with target depth of 10–50 million read pairs per sample. Library preparation blanks (no DNA template) were processed alongside samples to monitor for reagent contamination.

### Authentication of ancient DNA

To verify authenticity of ancient samples and distinguish genuine ancient DNA from modern contamination, we quantified characteristic postmortem DNA damage patterns using mapDamage v2.2.0 (*28*). Sequencing reads were first mapped to human reference genome GRCh38 using Bowtie2 v2.4.2 (*29*) with sensitive local alignment parameters (--very-sensitive-local). From mapped reads, mapDamage quantified cytosine-to-thymine (C→T) and guanine-to-adenine (G→A) substitution frequencies at 5′ and 3′ fragment termini, respectively, and determined fragment length distributions. Authentic ancient DNA exhibits elevated C→T substitutions at 5′ ends due to cytosine deamination during postmortem diagenesis. Samples were stratified by age and expected deamination patterns were verified.

### Quality control, contamination filtering, and dataset refinement

Raw sequencing reads were subjected to rigorous quality control and filtering. Illumina adapter sequences were trimmed using BBDuk2 from the BBMap package v38.96 (https://sourceforge.net/projects/bbmap/) with the following parameters: ktrim=r k=23 mink=11 hdist=1 tpe tbo. Low-quality bases (Phred score < 30) and reads shorter than 30 bp were removed using the same tool with parameters: qtrim=rl trimq=30 minlen=30. Read quality before and after filtering was assessed using FastQC v0.11.9 (https://www.bioinformatics.babraham.ac.uk/projects/fastqc/). Human DNA reads were depleted by mapping against the human reference genome GRCh38 using Bowtie2 v2.4.2 (*29*) with very-sensitive parameters (--very-sensitive), and unmapped reads (non-host microbial reads) were retained for downstream taxonomic and functional profiling.

### Taxonomic profiling

Taxonomic composition was profiled from the quality-filtered, non-host reads using the ucgMLST (universal core-genome Multi-Locus Sequence Typing) pipeline (*30*), which uses a reference database of universal single-copy core genes to provide high-resolution strain-level taxonomic assignments. Briefly, reads were mapped to the core gene database using minimap2 (*31*) mode with default parameters, and taxonomic assignments were made based on the best match with minimum alignment length of 30 nucleotides and minimum identity of 90%. Relative abundances were calculated by normalizing read counts to total microbial reads per sample and expressed as proportions. The resulting taxonomic profiles were compiled into a single abundance matrix with rows representing samples and columns representing species. The 20 most abundant species were identified based on their mean relative abundance across the entire dataset, accounting for 36.3% of total microbial abundance.

### Diversity analyses

All diversity and statistical analyses were performed in R v4.3.1 using the vegan v2.6-4 (https://cran.r-project.org/web/packages/vegan/) and phyloseq v1.44.0 packages (https://joey711.github.io/phyloseq/) unless otherwise specified. Alpha-diversity was calculated using the Simpson Diversity Index (SDI) on the species-level abundance data using the diversity function from vegan with method=“simpson”. The Simpson index ranges from 0 to 1, with higher values indicating greater diversity and evenness. Samples with SDI < 0.5 (n=54), predominantly from ancient dentin, were identified as having exceptionally low diversity indicative of severe DNA degradation, environmental contamination, or ecological collapse, and were excluded from further analyses. Beta-diversity was assessed using Bray-Curtis dissimilarity calculated from the species-level relative abundance matrix using the vegdist function with method=“bray”. Principal Coordinates Analysis (PCoA) was performed using the cmdscale function to reduce the dissimilarity matrix to two or three principal axes for visualization. The variance in community composition explained by anatomical site, temporal period, and geographic origin was quantified using Permutational Multivariate Analysis of Variance (PERMANOVA) implemented with the adonis2 function from vegan with 999 permutations. PERMANOVA tests with Benjamini-Hochberg FDR correction were performed to assess differences between specific groups. For visualization of the temporal and geographic structure within the dental calculus subset (n=606), Uniform Manifold Approximation and Projection (UMAP) (*32*)was implemented using the umap package v0.2.10.0 in R with the following parameters: n_neighbors=15, min_dist=0.1, metric=“euclidean”, applied to the centered log-ratio (CLR) transformed species abundance matrix.

### Machine learning classification for sample quality control and origin prediction

To objectively evaluate sample reliability and detect anatomical misclassification or contamination, we developed logistic (anatomical origin) or random forest classifiers (temporal and spatial origins) using the scikit-learn library v1.3.2 in Python v3.10.12 to predict the origin sof each sample from its microbial composition. The model used species-level relative abundances (310 species present in ≥5% of samples) as features and anatomical, temporal and spatial origins as the target variable. The dataset was randomly split into training (80%) and test (20%) sets with stratification to maintain class proportions. We used GridCV function with five-fold cross validation for searching hyperparameters for the models. Model performance was assessed using 5-fold stratified cross-validation on the training set, and accuracy and area under the receiver operating characteristic curve (AUC) were calculated on the held-out test set. Feature importance was determined using mean decrease in Gini impurity.

### Network analysis of co-occurrence modules

A species co-occurrence network was constructed from the dental calculus subset (n=606) to identify robust ecological modules. Species present in at least 5% of samples (n=310 species) were retained to ensure sufficient statistical power for correlation analysis. Relative abundances were transformed using the CLR method to account for the compositional nature of microbiome data and reduce spurious correlations. Pairwise Spearman rank correlations (ρ) were calculated for all species pairs using the cor.test function in R with method=“spearman”. To construct a sparse network retaining only robust associations, edges were retained if they satisfied two criteria: absolute correlation coefficient |ρ| > 0.6 and FDR-corrected p-value < 0.05 (Benjamini-Hochberg correction across all pairwise tests). The resulting network was visualized and analyzed using Gephi v0.10.1 (*33*). Microbial modules (communities) were identified using the Leiden algorithm, a modularity-optimization method implemented in Gephi, which iteratively assigns nodes to communities to maximize the modularity score Q.

The prevalence of each module within each sample was calculated as the summed relative abundance of all constituent species belonging to that module. Module prevalence values were then aggregated by temporal period and visualized as temporal trends. To quantify the ecological decoupling of DCM1 and DCM2 over time, inter-module and intra-module correlations were calculated for discrete temporal bins (>8,000 BP, 6,000–8,000 BP, 4,000–6,000 BP, 2,000–4,000 BP, 200–2,000 BP, <200 BP). For each bin, all pairwise Spearman correlations were calculated between species, and correlations were classified as intra-module (both species in same module) or inter-module (species in different modules). The mean correlation coefficient was calculated for each category within each temporal bin, and the difference (Δ) between intra– and inter-module correlation was computed as a metric of ecological polarization.

### Temporal Trend Analysis

To quantify individual species’ contributions to long-term temporal shifts and assess whether trends varied by geography, we modeled associations between microbial relative abundances and sample age using generalized linear models (GLMs). For each of 310 species present in at least 15% of dental calculus samples, we fitted a GLM with quasi-Poisson distribution to accommodate overdispersion in count-based abundance data. The model formula was: log (abundance + pseudocount) ∼ log(age_BP) + geographic_region, where abundance represents the species relative abundance multiplied by 10[ and rounded to integers, pseudocount was set to 1 to handle zeros, age_BP is sample age in years before present, and geographic_region is a categorical variable with levels Europe, Asia, Americas, and Africa. The quasi-Poisson family with log link was chosen to model non-negative, overdispersed count data.

Statistical significance of the age coefficient (temporal trend) was assessed using likelihood ratio tests comparing full models to reduced models without the age term, with FDR correction (Benjamini-Hochberg, q<0.05) across all 310 species. This analysis identified significant temporal dynamics in 145 of 310 species (46.8%), indicating widespread temporal shifts in the taxonomic composition of the oral microbiome (**Figure 3a**). Species with significant temporal trends were then classified into “increasing” or “decreasing” groups based on the sign and magnitude of their age coefficient using Gaussian Mixture Models (GMM). GMM was implemented using the sklearn.mixture.GaussianMixture function in Python v3.10 with n_components=2, covariance_type=’full’, and 100 random initializations. The model was fitted to the vector of age coefficients from significant species, and classification into two clusters yielded 63 taxa classified as increasing and 82 as decreasing in abundance over time (**Figure 3c**). To assess geographic variation in temporal trends, we repeated the GLM analysis stratified by geographic region (Europe, Asia, Americas, Africa separately), comparing the magnitude and direction of age coefficients across regions using Kruskal-Wallis tests. Regional stratification revealed that European populations exhibited more pronounced and consistent temporal changes compared to African and American populations.

### Phylogenetic reconstruction

For two key species with contrasting evolutionary trajectories (*Pauljensenia mediterranea* and *Actinomyces israelii*), we performed detailed phylogenetic reconstruction and population structure analysis. For each species, metagenomic reads were aligned to a reference genome using EToKi assemble function (*34*), which internally uses minimap2 (*31*) for alignment and reconstructs consensus base calls of the core genome using samtools (*35*). Only genomes with at least 50% of its core genome covered by 3× or more reads were retained. All reconstructed genomes were aligned to the reference genomes using EToKi align and had the maximum-likelihood phylogenetic trees constructed using IQ-TREE v2.2.0 (*36*).

### Temporal breakpoint analysis for detecting cultural transition signatures

To pinpoint specific periods of accelerated community-wide ecological change corresponding to major cultural transitions, we performed a multivariate breakpoint analysis on time-ordered species abundance data. The analysis was designed to detect significant shifts in the overall microbial community structure rather than individual species trends. We applied a dynamic programming change-point detection algorithm implemented in the ruptures Python library v1.1.8 (*37*), specifically using the Pelt (Pruned Exact Linear Time) method with an L2 cost function (squared Euclidean distance) to efficiently identify multiple breakpoints in multivariate time series. The input data consisted of the CLR-transformed species abundance matrix from dental calculus samples (n=606, 310 species), ordered chronologically by sample age. Prior to analysis, species abundances were standardized (z-score normalized) to ensure equal weighting across taxa. The Pelt algorithm was run with a penalty parameter β=10 to control the number of detected breakpoints, chosen based on cross-validation to balance sensitivity and specificity. To assess statistical significance, we performed permutation testing by randomly shuffling sample ages 1,000 times and recalculating breakpoint densities, defining significant breakpoints as those exceeding the 95th percentile of the null distribution.

### Functional and metabolic Analysis

Functional potential of the oral microbiome was characterized by profiling gene content and reconstructing metabolic pathways from the quality-filtered, non-host metagenomic reads. Functional annotation was performed using HUMAnN 3.0 (The HMP Unified Metabolic Analysis Network) (*38*). Pathway abundances were mapped to KEGG orthology (KO) terms and KEGG pathways using the HUMAnN utility humann_regroup_table with the KEGG mapping files (v2023.1). Only pathways with >1 CPM abundance in at least 15% of dental calculus samples were retained for temporal analysis. To identify pathways and genes with significant temporal trends, we applied the same GLM framework used for taxonomic data: log (pathway_abundance + 1) ∼ log(age_BP) + geographic_region, using quasi-Poisson family and FDR correction (q<0.05).

### Antimicrobial resistance gene profiling and classification

To characterize the evolution of antimicrobial resistance in the oral microbiome and assess the impact of antibiotic use on resistome composition, we profiled antimicrobial resistance genes (ARGs) from quality-filtered metagenomic reads using the ARGs-OAP v3.0 pipeline (Antibiotic Resistance Gene-Online Analysis Pipeline) (*39*). ARG abundance was quantified as the number of reads mapping to each ARG normalized by gene length and total sequencing depth, expressed as reads per kilobase per million mapped reads (RPKM): RPKM = (number of reads mapped to ARG × 10[) / (ARG length in bp × total non-host reads). Only ARGs detected with ≥95% sequence identity over ≥80% alignment length was retained to ensure annotation accuracy. ARGs were classified into functional categories based on antibiotic target class (beta-lactams, tetracyclines, macrolides, aminoglycosides, etc.) and mechanism of resistance (efflux pumps, target modification, antibiotic inactivation, target protection).

To assess epidemiological and evolutionary implications, ARGs were further stratified into four mobility ranks (I–IV) according to the SARG database classification framework (*14*), which reflects clinical relevance and horizontal gene transfer (HGT) potential: Rank I includes highly mobile, clinically urgent ARGs frequently located on plasmids, transposons, or integrons and associated with current clinical treatment failures (e.g., carbapenemases, ESBL genes, vancomycin resistance genes); Rank II includes mobile ARGs with moderate clinical relevance, often found on mobile genetic elements but less frequently reported in clinical settings; Rank III includes ARGs with low mobility, typically chromosomal, representing intrinsic or species-specific resistance mechanisms; Rank IV includes putative ARGs identified by sequence homology but lacking functional validation or known clinical significance. This hierarchical classification enables discrimination between stable, background resistome components and rapidly evolving, clinically concerning resistance determinants.

To identify key microbial taxa harboring and potentially disseminating resistance determinants, we constructed ARG–host association networks based on co-abundance patterns across samples. For each ARG detected in ≥5% of samples and each species detected in ≥5% of samples, Spearman rank correlations were calculated between ARG abundance (RPKM) and species relative abundance using CLR-transformed data to account for compositionality. Significant associations (|ρ| > 0.3, FDR-corrected p < 0.05) were retained as edges in a bipartite network with ARGs and species as node types.

### Regional connectivity analysis

To assess regional patterns of oral microbiome similarity and identify geographic clusters or isolates, we constructed sample-to-sample similarity networks based on taxonomic composition. Pairwise Spearman rank correlations (ρ) were calculated between all dental calculus samples (n=606) using CLR-transformed species abundance profiles. A network was constructed by retaining edges (sample pairs) with correlation coefficients ρ > 0.6 and FDR-corrected p-values < 0.05, thresholds chosen to identify robust similarities while maintaining computational tractability. The resulting network was visualized in Gephi v0.10.1, with node sizes scaled by degree centrality and colors coded by geographic region. Network topology was quantified using several centrality measures: degree centrality (number of connections per node), eigenvector centrality (influence based on connections to other highly connected nodes), and betweenness centrality (frequency of lying on shortest paths between other nodes). Regional connectivity was assessed by calculating within-region and between-region edge densities and comparing these to null expectations from 1,000 permutations of region labels.

### Hierarchical clustering

To further elucidate the hierarchical relationships and relative contributions of spatial versus temporal drivers of microbial community variation, we performed hierarchical clustering on population-level taxonomic centroids. Samples were first grouped into spatiotemporal bins defined by the combination of geographic region (Europe, Asia, Americas, Africa) and temporal period (0–100 years BP, 100–200 years BP, 200–500 years BP, 500–1,000 years BP, 1,000–2,000 years BP, 2,000–4,000 years BP, 4,000–8,000 years BP, >8,000 years BP). For each bin, a regional centroid was computed as the mean relative abundance vector across all constituent samples. Pairwise Bray-Curtis dissimilarities were calculated in python. Hierarchical clustering was performed on the dissimilarity matrix online tools in SRPlot (*14*).

### Statistical analyses

Unless otherwise noted, all statistical tests were two-sided and performed in R v4.3.1. Multiple hypothesis testing correction was applied using the Benjamini-Hochberg false discovery rate (FDR) method with q < 0.05 as the significance threshold unless otherwise specified. For continuous variables, normality was assessed using Shapiro-Wilk tests, and non-parametric tests (Wilcoxon rank-sum, Kruskal-Wallis) were used when distributions significantly deviated from normality (*p* < 0.05). For categorical comparisons, Fisher’s exact test or chi-square test were used depending on expected cell counts. Effect sizes were reported as Cohen’s d for t-tests, Cliff’s delta for rank-based tests, and η² for ANOVA/PERMANOVA tests. Confidence intervals (95% CI) were calculated using bootstrap resampling with 1,000 iterations where analytical solutions were unavailable.

## Supporting information

The numbers of samples enrolled in this study.

Detail information of the samples enrolled in this study.

Information of species and eignvector centrality in two modules.

Regional stratification analysis.

KEGG pathway annotations and KOs down-or up-regulated in dental calculus microbiome.

## ACKNOWLEDGMENTS

We thank Sandra Bedarida for helps in handling ancient materials; Grégory Pereira for kindly offering Mexican samples; Daniel Falush, Xin Lu, and Ping He for their support and insight; and Iotabiome Biotechnology Inc. for their technical support. The project was primarily supported by Wellcome Trust (202792) and the Shenzhen Medical Research Fund (B2403009). This work was also supported by the National Natural Science Foundation of China (82530102, 32170003, 32370099), the Provincial-level Talent Program for National Center of Technology Innovation for Biopharmaceuticals (NCTIB2024JS0101), the Natural Science Foundation of Jiangsu Province (BK20243008), and the Suzhou Top-Notch Talent Groups (ZXD2022003).

## AUTHOR CONTRIBUTIONS

M.A. and Z.Z. conceptualized the project and designed the study. R.B., S.S., C.S., L.S.W. contributed study materials. L.Z., H.L., S.X., S.X., S.L., Q.S., and H.Y. analyzed and interpreted data. L.Z. and Z.Z. visualized the results and prepared figures. L.Z., H.L., and Z.Z. wrote the initial draft of the manuscript. M.A., H.L., S.X., Z.Z., Y.C, and H.Y. discussed and revised the manuscript. All authors edited and commented on the manuscript.

## Supplementary Figure Legends

**Supplementary Figure 1.**
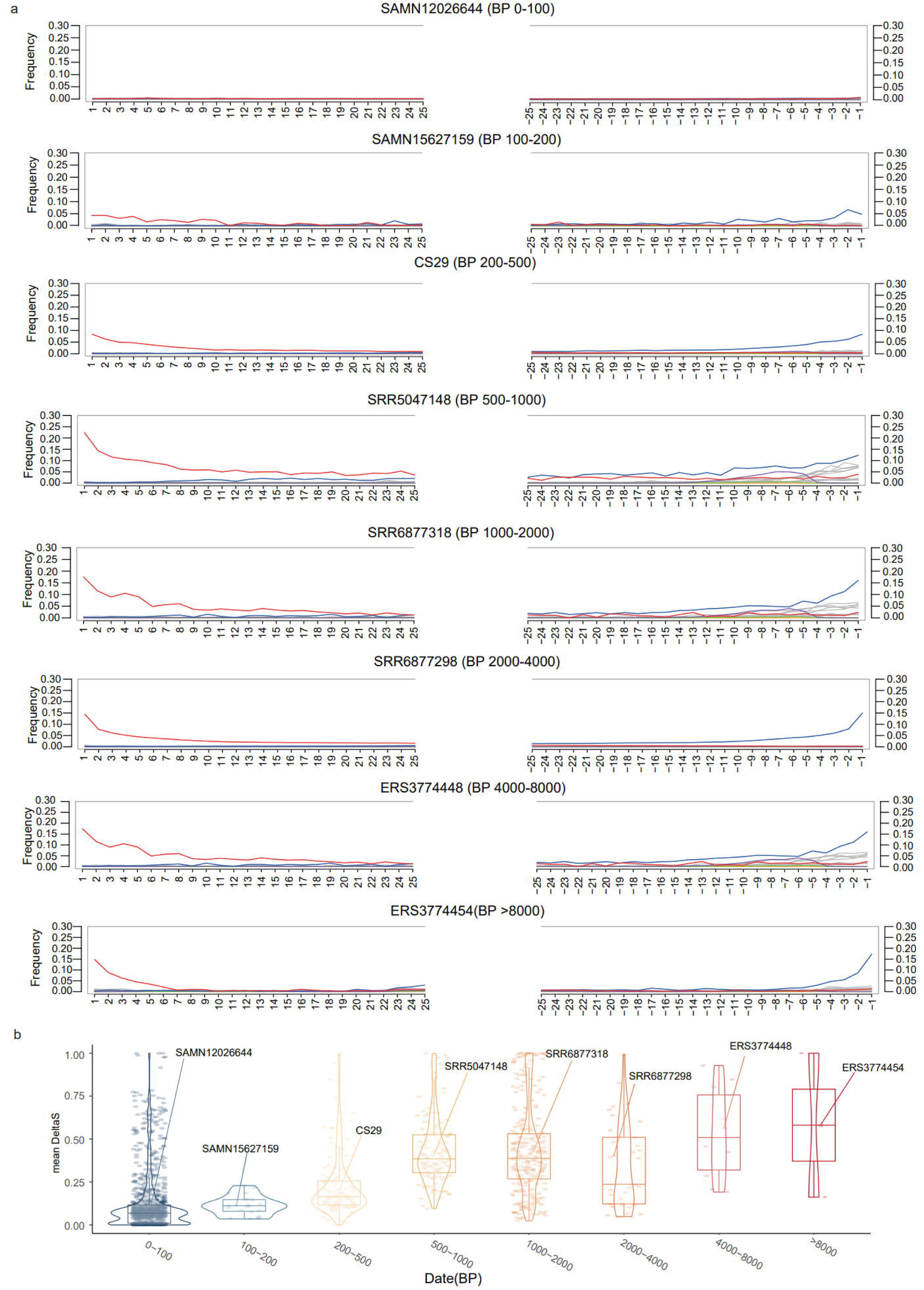
Post-mortem DNA damage patterns across temporal bins validate ancient DNA authenticity. **(a)** Frequency of cytosine-to-thymine (C→T) substitutions at the 5′ ends of DNA fragments for representative samples across eight temporal bins (0-100, 100-200, 200-500, 500-1000, 1000-2000, 2000-4000, 4000-8000, and >8000 years BP). Each panel shows substitution frequencies at positions relative to fragment termini (x-axis: 1-25 bases from 5′ end on left, –25 to –1 bases from 3′ end on right). Red lines indicate C→T misincorporation rates characteristic of post-mortem cytosine deamination. Older samples (bottom panels) display elevated C→T substitution frequencies at terminal positions. Sample identifiers and temporal bins are indicated above each panel. **(b)** Violin plots summarizing the distribution of DNA damage levels across all samples grouped by temporal bin. Y-axis shows mean damage (0-1.00 scale representing 0-100% C→T substitution rate at terminal positions). Box plots within each violin indicate median (horizontal line), interquartile range (box), and range (whiskers). Gray points on the leftmost violin represent individual modern samples. Representative samples from panel (a) are labeled above their respective distributions.

**Supplementary Figure 2.**
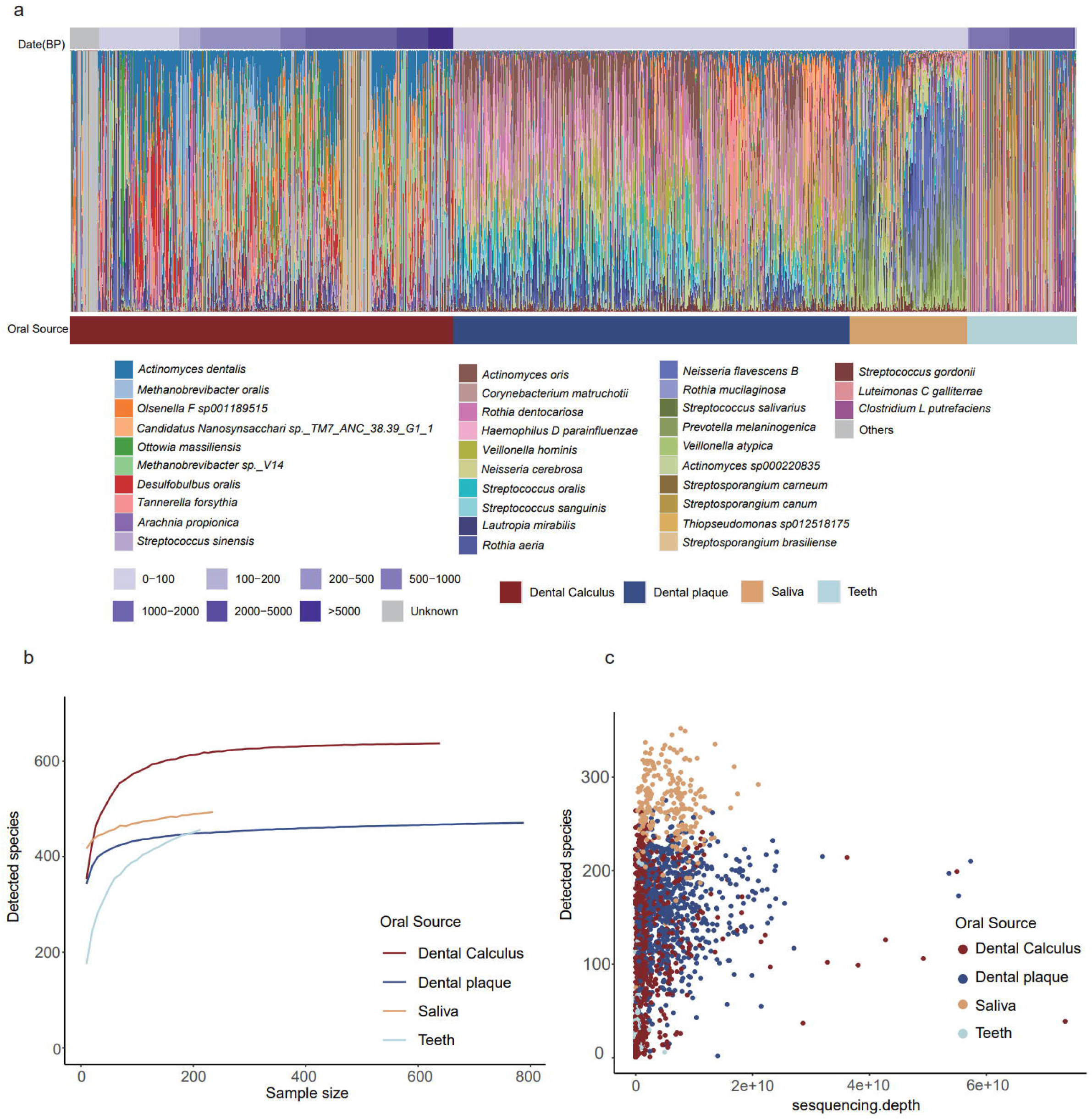
Taxonomic composition and species richness across oral sources and temporal periods. **(a)** Stacked bar plot showing the taxonomic composition of oral microbiomes across individual samples. Each vertical bar represents a single sample, with colors indicating different microbial species. The top 20 most abundant species (comprising 36.3% of total microbial abundance) are individually colored and labeled in the legend, while remaining taxa are grouped as “Others” (gray). Samples are organized by oral source (bottom annotation bar): dental calculus (dark red), dental plaque (dark blue), saliva (tan), and teeth (light gray). The temporal age of samples is indicated by the color gradient in the top annotation bar, ranging from modern (0-100 BP, light purple) to ancient (>5000 BP, dark purple). **(b)** Rarefaction curves showing the relationship between sample size (x-axis, number of reads) and detected species richness (y-axis, number of species) for each oral source. Curves are color-coded by source: dental calculus (dark red), dental plaque (dark blue), saliva (tan), and teeth (light gray). **(c)** Scatter plot showing the relationship between sequencing depth (x-axis, total sequencing reads per sample, displayed in scientific notation) and detected species richness (y-axis, number of species) for individual samples. Each point represents a single sample, colored by oral source using the same scheme as panel (b).

**Supplementary Figure 3.**
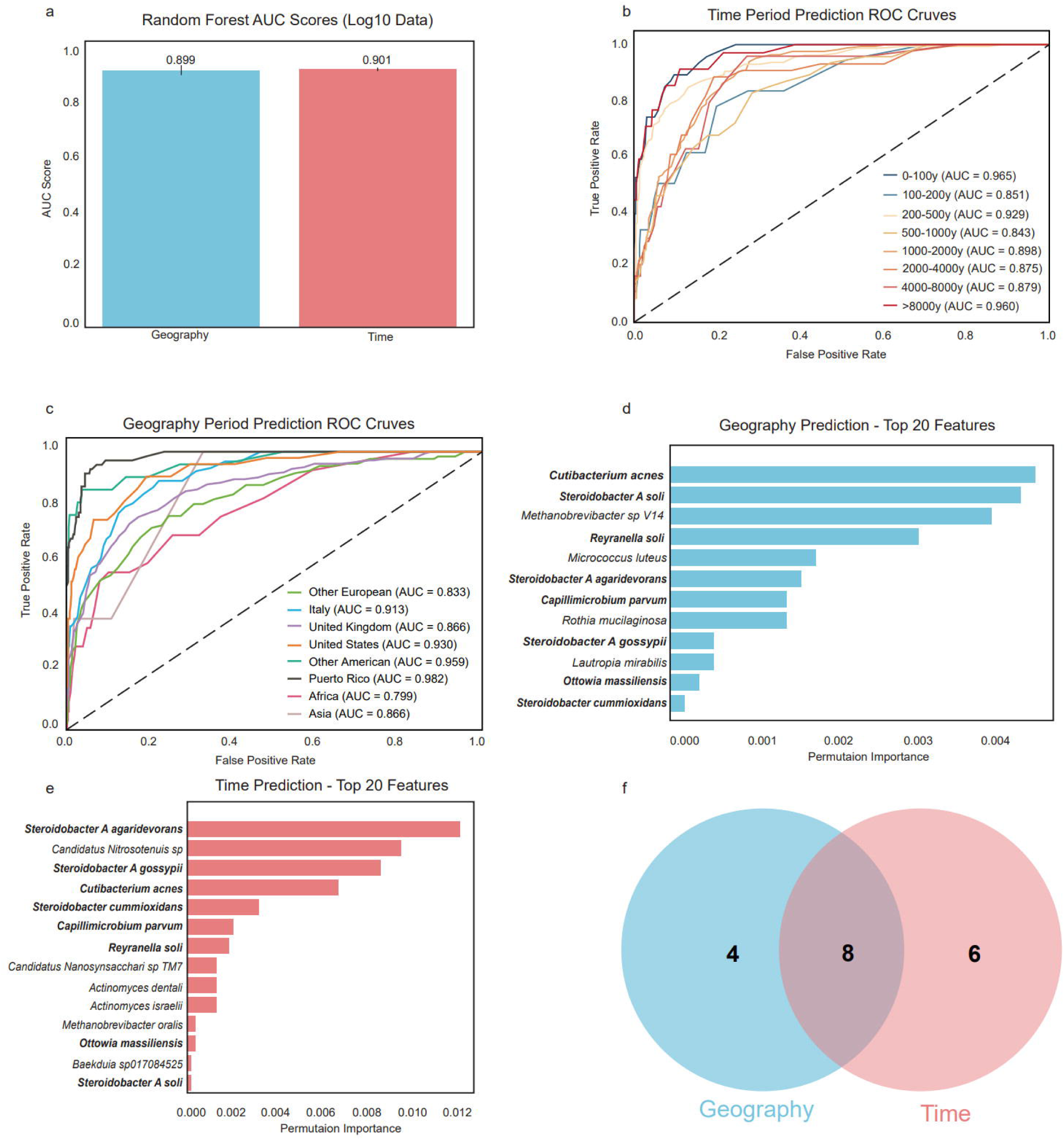
Machine learning classification of oral microbiomes by geography and time reveals distinct taxonomic signatures. **(a)** Random forest model performance for predicting sample origin based on taxonomic profiles using log10-transformed abundance data. Bar plot shows Area Under the Curve (AUC) scores for geographic region prediction (blue, AUC = 0.899) and temporal period prediction (pink, AUC = 0.901). **(b)** Receiver Operating Characteristic (ROC) curves for geography prediction across different geographic regions. Each colored line represents a one-vs-rest classification for a specific region, with AUC scores shown in the legend. The diagonal dashed line represents random chance (AUC = 0.5). **(c)** ROC curves for temporal period prediction across eight time bins. **(d)** Twelve significant microbial features ranked by permutation importance for geography prediction. Horizontal bars indicate the relative contribution of each taxon to model accuracy, with longer bars representing greater predictive importance. **(e)** Fourteen significant microbial features ranked by permutation importance for temporal period prediction. **(f)** Venn diagram illustrating the overlap between geographically informative taxa (blue circle, 4 unique taxa) and temporally informative taxa (pink circle, 6 unique taxa). Eight taxa (central overlap) contribute to both geographic and temporal predictions, representing 67% of geographic predictors and 57% of temporal predictors.

**Supplementary Figure 4.**
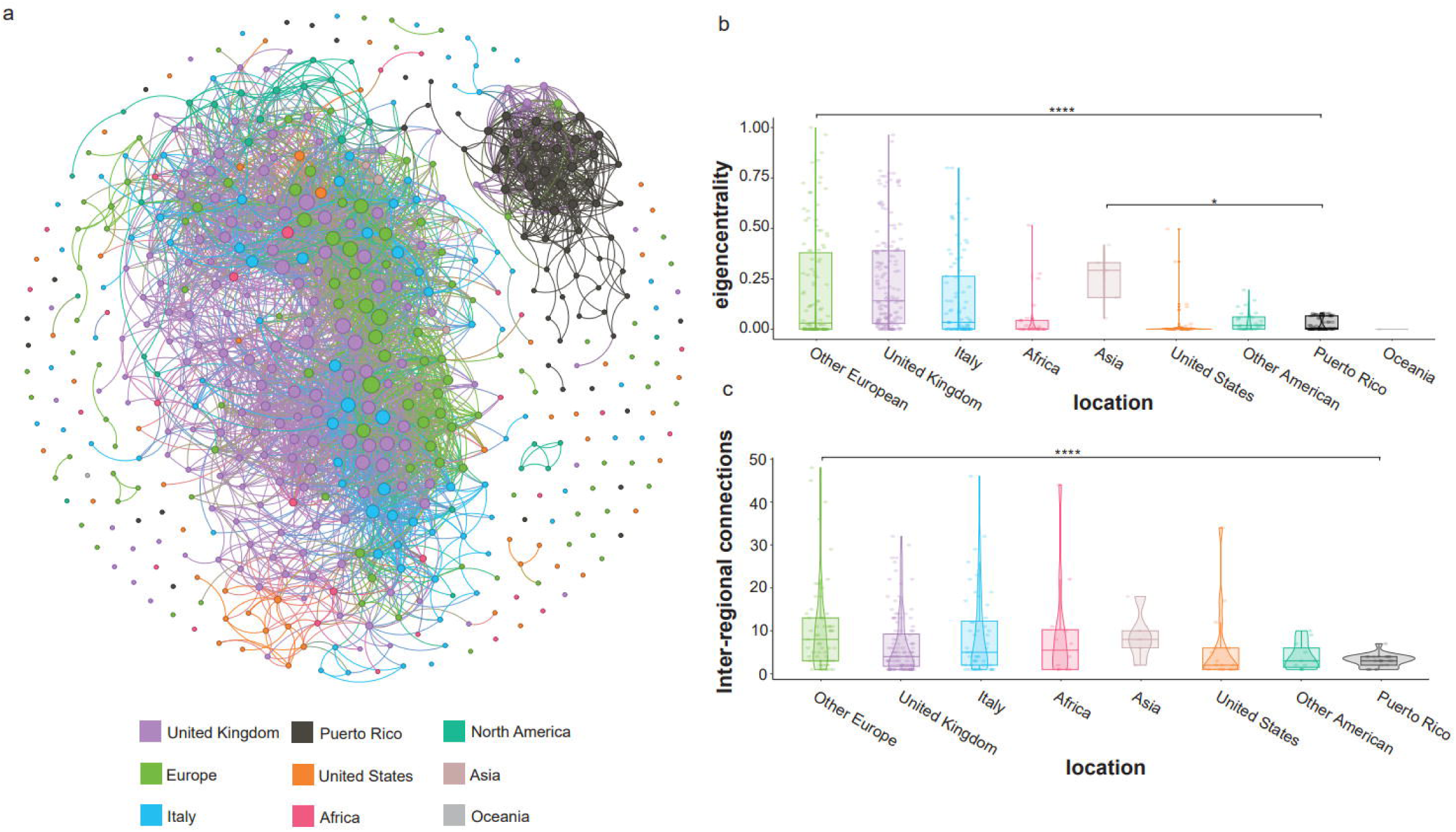
Network analysis reveals geographic patterns of oral microbiome connectivity and regional isolation. **(a)** Network visualization of sample-to-sample relationships based on Spearman’s correlation coefficients of taxonomic profiles. Each node represents an individual sample, with node size proportional to the number of connections (degree centrality). Edges (lines) connect samples with significant positive correlations in microbial composition, with edge density reflecting the strength of microbial similarity. Nodes are colored by geographic region according to the legend. **(b)** Eigenvector centrality distributions across geographic regions. **(c)** Inter-regional connection counts across geographic regions. This metric quantifies the average number of connections each sample has to samples from different geographic regions, reflecting cross-regional microbiome similarity and potential historical population contact.

**Supplementary Figure 5.**
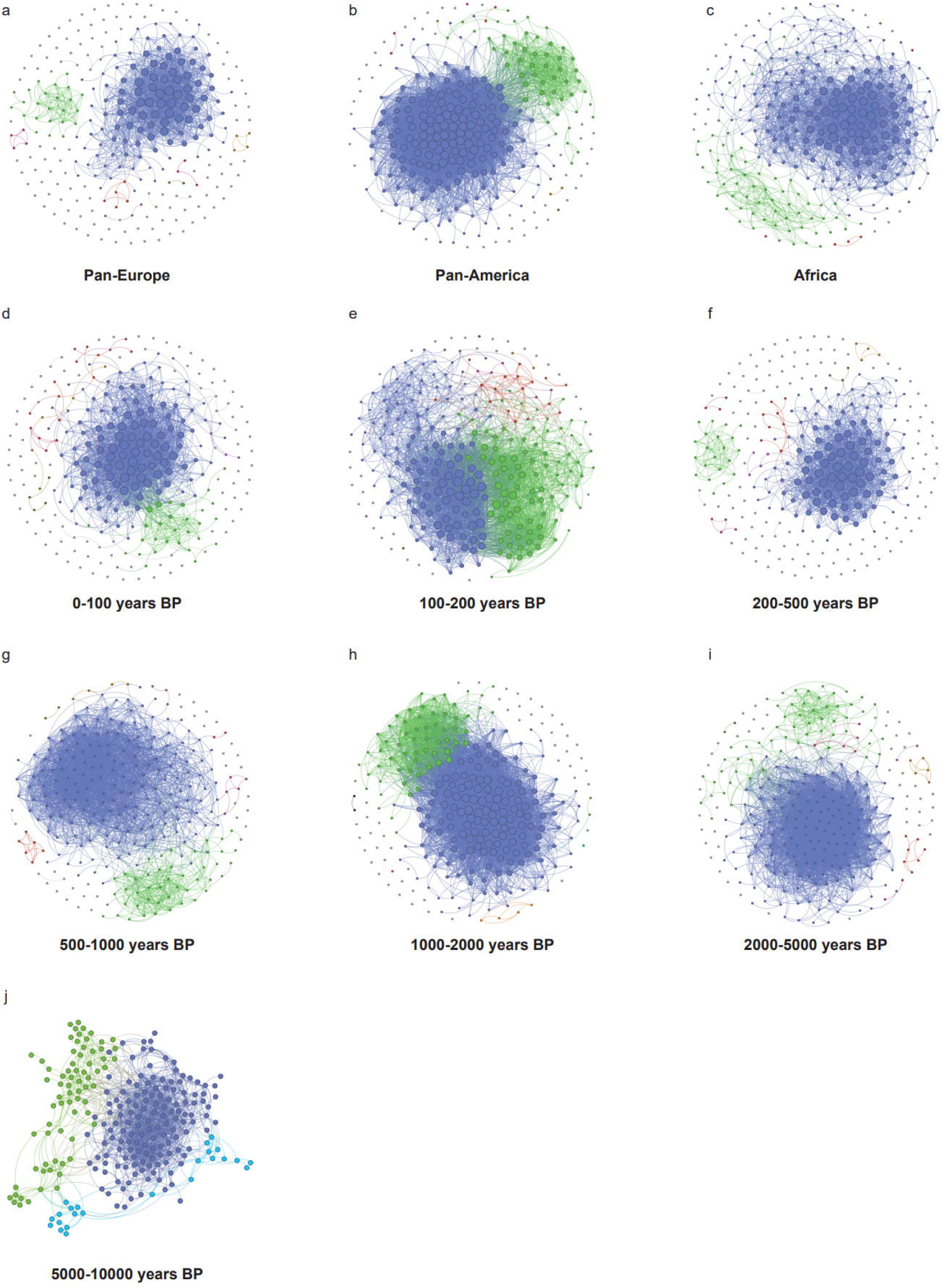
Network structures of species co-occurrence patterns across geographic regions and temporal bins. Force-directed ecological networks depicting pairwise co-occurrence relationships among oral microbial species in dental calculus samples, stratified by **(a–c)** major geographic regions and **(d–j)** calibrated temporal intervals (BP, before present). Each node represents a microbial species, and edges indicate significant positive co-occurrence relationships. Node colors correspond to community modules inferred from network clustering, highlighting the two major dental calculus modules (DCM1 and DCM2). The spatial panels show networks from (a) Pan-Europe, (b) Pan-America, and (c) Africa. Temporal panels include samples dated to (d) 0–100 years BP, (e) 100–200 years BP, (f) 200–500 years BP, (g) 500–1000 years BP, (h) 1000–2000 years BP, (i) 2000–5000 years BP, and (j) 5000–10,000 years BP.

**Supplementary Figure 6.**
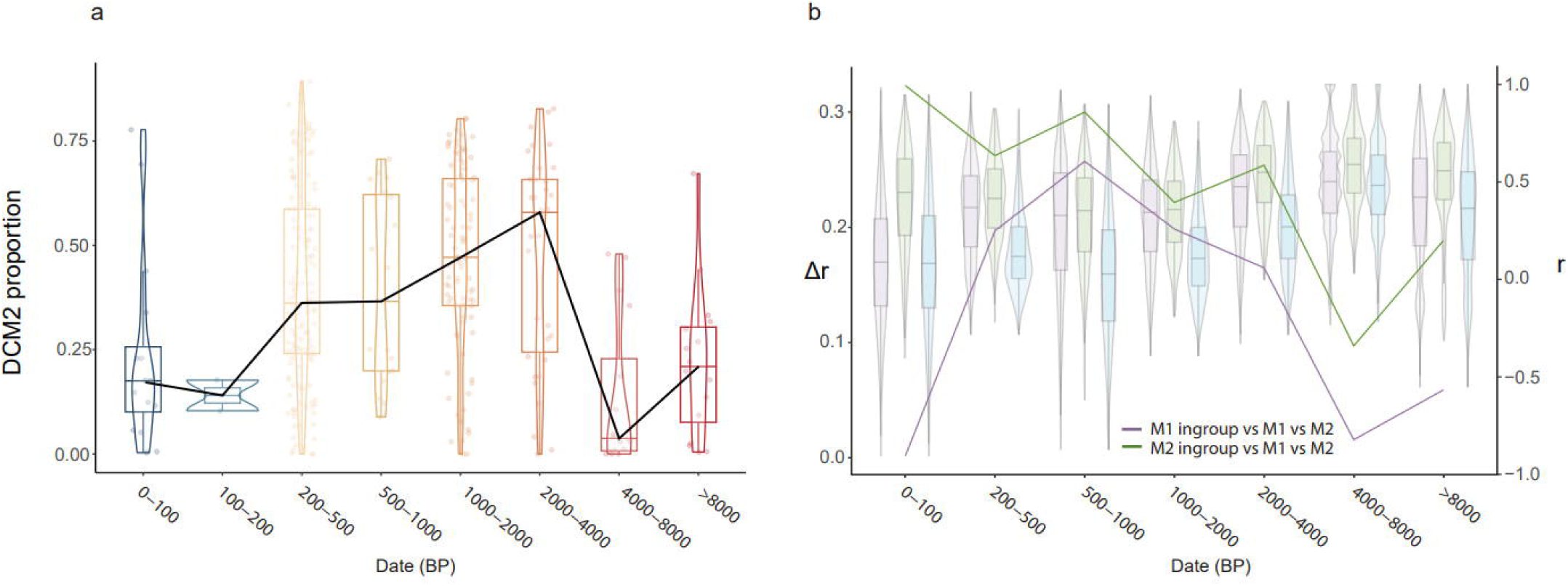
Temporal dynamics of pathogen-associated module prevalence and long-term ecological decoupling between oral microbiome modules. **(a)** Changes in the proportional abundance of the pathogen-associated dental calculus module (DCM2) across calibrated temporal intervals (BP, before present). Violin plots show the distribution of individual sample proportions within each time bin, with overlaid box plots summarizing medians and interquartile ranges. Superimposed line traces indicate the median DCM2 proportion through time. **(b)** Inter-module correlation structure across the same temporal bins. Violin and box plots display the distribution of pairwise species-level correlations (r) within DCM1 (commensal module) and DCM2 (pathogen-associated module), as well as between modules. Overlaid lines depict the temporal trajectory of correlation differences (Δr = intra-module r – inter-module r) for DCM1 (purple) and DCM2 (green).

**Supplementary Figure 7.**
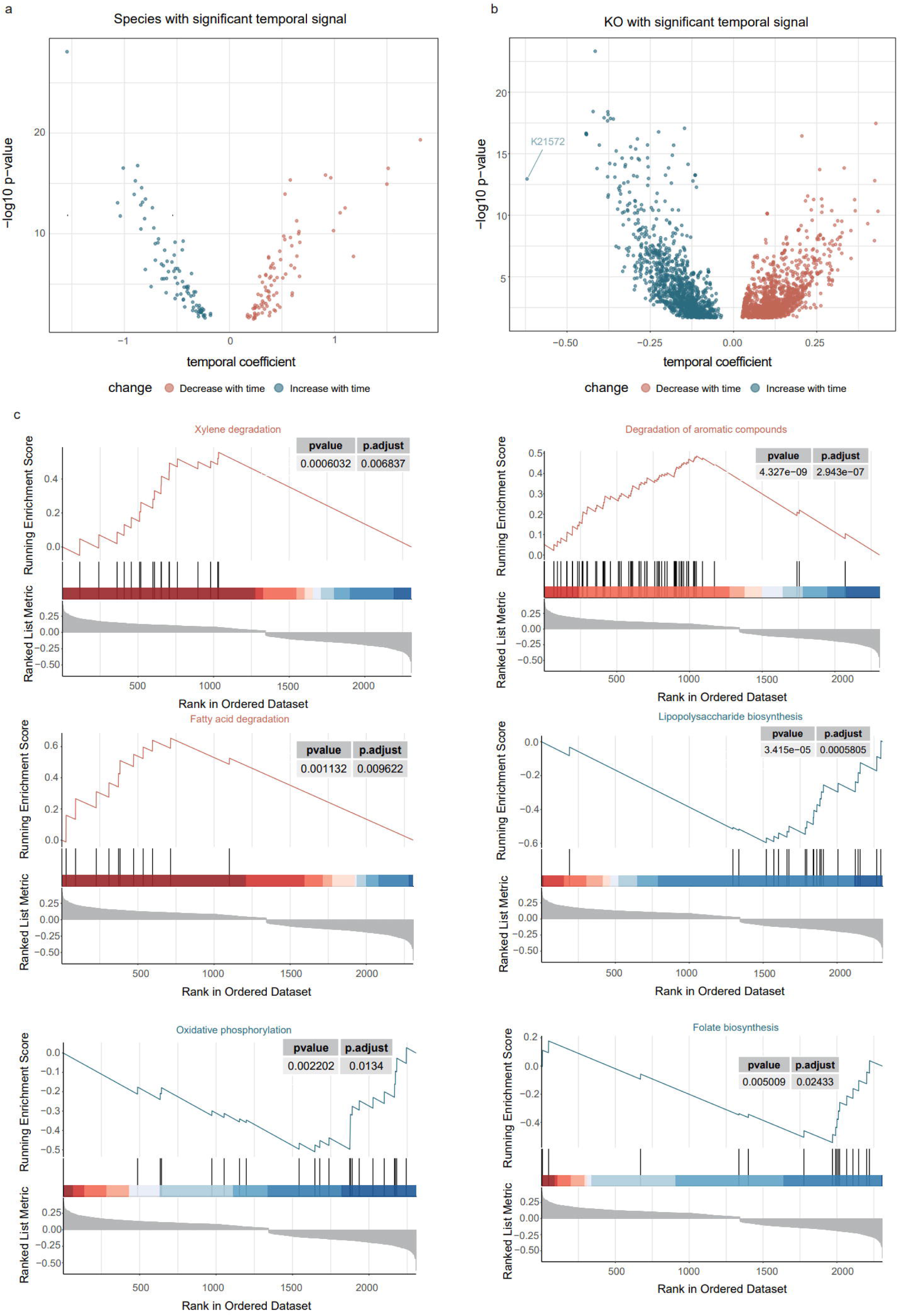
Taxonomic and functional signatures of long-term temporal change in the oral microbiome. **(a)** Volcano plot showing microbial species with significant temporal associations, estimated using generalized linear models with sample age as a continuous predictor and geographic origin as a covariate. Points represent species, plotted by temporal coefficient (x-axis) and –log10 *P* value (y-axis). Species with significantly increasing abundance through time are shown in blue, and those decreasing in red. **(b)** Temporal signal among KEGG Orthologs (KOs), highlighting functional genes whose abundances exhibit significant temporal trends. Each point represents a KO; colors denote increasing (blue) or decreasing (red) temporal trajectories. **(c)** Gene Set Enrichment Analysis (GSEA) of temporally ordered KOs, identifying metabolic pathways enriched among genes increasing (red) or decreasing (blue) through time. Example pathways with significant enrichment include *xylene degradation*, *degradation of aromatic compounds*, and *fatty acid degradation* (decreasing with time), as well as *lipopolysaccharide biosynthesis*, *oxidative phosphorylation*, and *folate biosynthesis* (increasing with time). For each pathway, running enrichment scores and ranked KO metrics are shown alongside nominal *P* values and adjusted *P* values.

**Supplementary Figure 8.**
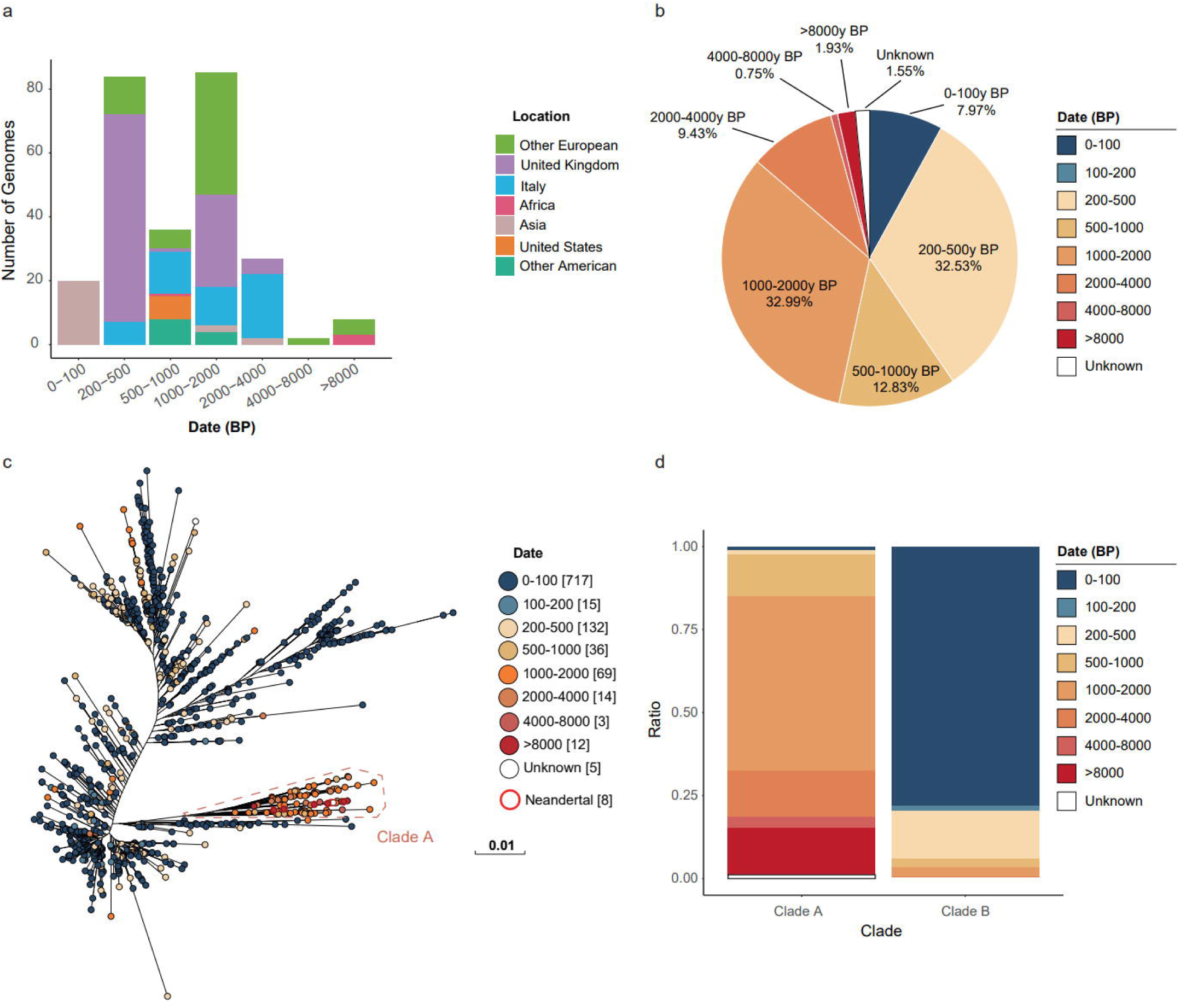
Temporal population dynamics and phylogenetic restructuring of two oral microbiome species with the strongest long-term temporal signals. **(a)** Temporal distribution of *Pauljensenia mediterranea* genomes across geographic regions, showing a dramatic contraction in population size and spatial range over the past 200 years. **(b)** Proportion of genomes assigned to each temporal bin, highlighting the overwhelming dominance of ancient samples and the near disappearance of the species in recent centuries. **(c)** Maximum-likelihood phylogeny of *Actinomyces israelii*, colored by sample age, revealing two deeply divergent lineages (Clades A and B). Clade A comprises predominantly ancient genomes spanning 208–102,400 BP, including several Neanderthal individuals, and displays a historically widespread distribution across Eurasia. **(d)** Relative abundance of temporal bins within Clades A and B, demonstrating a pronounced clade replacement event beginning ∼1000 BP: Clade A sharply declines and becomes absent in the past 200 years, whereas Clade B, rare in samples older than 2000 BP, expands rapidly and dominates in the last millennium.

**Supplementary Figure 9.**
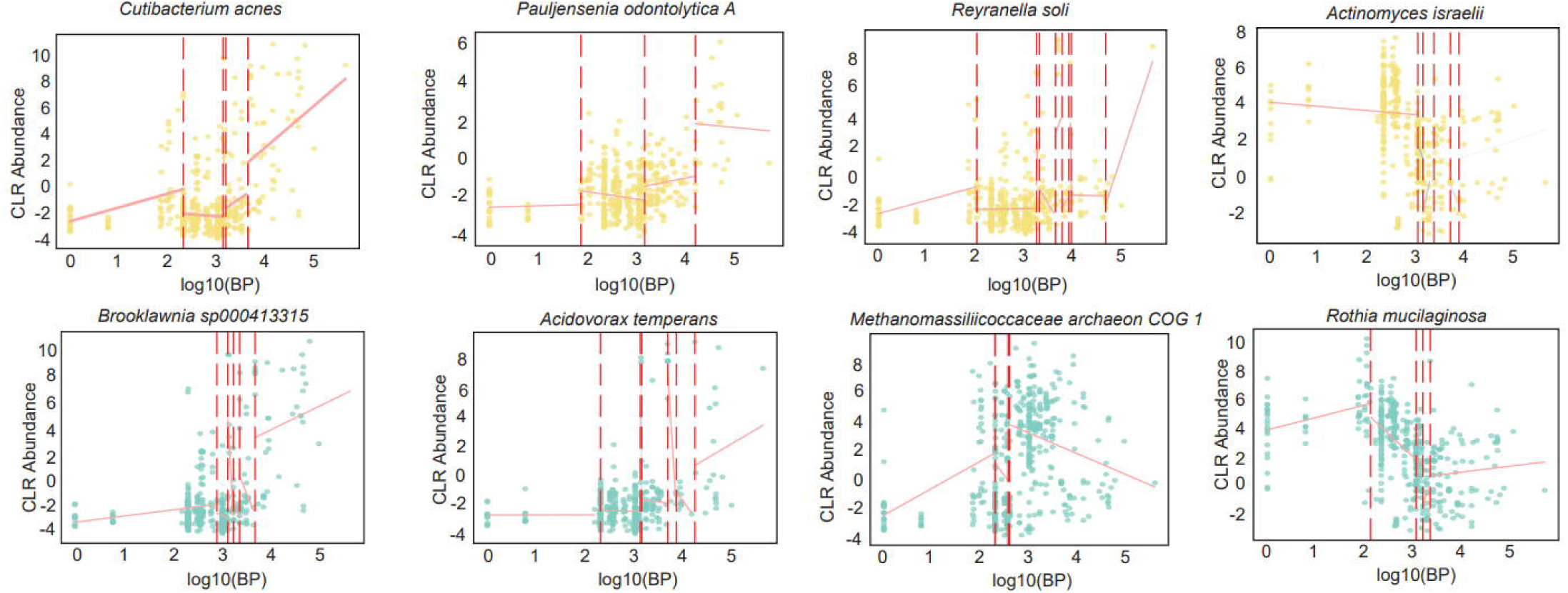
Species-level breakpoint analysis reveals temporal shifts in oral microbial abundance trajectories coincident with major cultural transitions. Each panel displays the CLR-transformed abundance (y-axis) of a representative oral taxon against time (x-axis, log[[(BP)) for European samples. Red dashed vertical lines indicate statistically significant breakpoints identified via changepoint analysis; clusters around ∼3.0 (5000 BP) and ∼2.3 (200 BP) correspond to the Neolithic and Industrial Revolutions, respectively. A smoothed trend line (pink) illustrates overall abundance dynamics. Breakpoints are concentrated during these pivotal cultural periods across multiple taxa, suggesting that major societal transitions drove accelerated restructuring of the human oral microbiome. Taxa shown include *Cutibacterium acnes*, *Pauljensenia odontolytica A*, *Reyrannella soli*, *Actinomyces israelii*, *Brooklawnia sp000413315*, *Acidovorax temperans*, *Methanomassiliicoccaceae archaeon COG 1*, and *Rothia mucilaginosa*.

