## Supplementary material for "Evolutionary landscape of oral microbiome over 100,000 years": The numbers of samples enrolled in this study.

**Supplementary Table 1. The numbers of samples enrolled in this study.**

| **Oral Source** | Numbers of samples |
| --- | --- |
| Dental Calculus | 617 |
| Dental plaque | 788 |
| Saliva | 235 |
| Teeth | 214 |
| **Continent** |  |
| Africa | 33 |
| Asia | 268 |
| Europe | 602 |
| Americas | 928 |
| Oceania | 23 |
| **Country** |  |
| United States | 789 |
| China | 185 |
| United Kingdom | 174 |
| Poland | 167 |
| Puerto Rico | 110 |
| Italy | 95 |
| Ireland | 71 |
| Thailand | 46 |
| Spain | 38 |
| Philippines | 27 |
| Others | 152 |
| **Date (BP) group** |  |
| 0-100 | 1073 |
| 100-200 | 22 |
| 200-500 | 162 |
| 500-1000 | 96 |
| 1000-2000 | 340 |
| 2000-4000 | 47 |
| 4000-8000 | 24 |
| >8000 | 39 |
| Unknown | 51 |
| **Host species** |  |
| *Homo sapiens* | 1839 |
| *Homo sapiens neanderthalensis* | 15 |
